# Computational Mapping of the Human-SARS-CoV-2 Protein-RNA Interactome

**DOI:** 10.1101/2021.12.22.472458

**Authors:** Marc Horlacher, Svitlana Oleshko, Yue Hu, Mahsa Ghanbari, Giulia Cantini, Patrick Schinke, Ernesto Elorduy Vergara, Florian Bittner, Nikola S. Mueller, Uwe Ohler, Lambert Moyon, Annalisa Marsico

**Affiliations:** Computational Health Center, Helmholtz Center Munich, Germany; Berlin Institute for Medical Systems Biology, Max Delbrück Center for Molecular Medicine, Berlin, Germany; Knowing01 GmbH, Munich, Germany

**Keywords:** SARS-CoV-2, RBP binding, deep learning

## Abstract

Strong evidence suggests that human human RNA-binding proteins (RBPs) are critical factors for viral infection, yet there is no feasible experimental approach to map exact binding sites of RBPs across the SARS-CoV-2 genome systematically at a large scale. We investigated the role of RBPs in the context of SARS-CoV-2 by constructing the first in silico map of human RBP / viral RNA interactions at nucleotide-resolution using two deep learning methods (pysster and DeepRiPe) trained on data from CLIP-seq experiments. We evaluated conservation of RBP binding between 6 other human pathogenic coronaviruses and identified sites of conserved and differential binding in the UTRs of SARS-CoV-1, SARS-CoV-2 and MERS. We scored the impact of variants from 11 viral strains on protein-RNA interaction, identifying a set of gain-and loss of binding events. Lastly, we linked RBPs to functional data and OMICs from other studies, and identified MBNL1, FTO and FXR2 as potential clinical biomarkers. Our results contribute towards a deeper understanding of how viruses hijack host cellular pathways and are available through a comprehensive online resource (https://sc2rbpmap.helmholtz-muenchen.de).

## 1 Introduction

SARS-CoV-2, causative agent of the recent COVID-19 pandemic, has and still is affecting the lives of billions of people worldwide. Despite the large-scale vaccination effort, the number of infections and deaths remains high, primarily among the non-vaccinated and otherwise vulnerable individuals. Difficulty to control SARS-CoV-2 infections is partly due to the continuous emergence of novel viral variants, against which the full efficacy of current vaccines is still debated, as well as the lack of effective medication. This calls for a better understanding of the biology of SARS-CoV-2 to design alternative therapeutic strategies. SARS-CoV-2 is a betacoronavirus with a positive-sense, single-stranded RNA of ~30kb (90). Upon infection, the released RNA molecule depends on the host cell’s protein synthesis machinery to express a set of viral proteins crucial for replication (73). The genomic RNA is translated to produce non-structural proteins (nsps) from two open reading frames (ORFs), ORF1a and ORF1b, and it also contains untranslated regions (UTRs) at the 5’ and 3’ ends of the genomic RNA (90). A recent study revealed the complexity of the SARS-CoV-2 transcriptome, due to numerous discontinuous transcription events (39). Negative sense RNA intermediates are generated to serve as the template for the synthesis of positive-sense genomic RNA (gRNA) and subgenomic RNAs (sgRNA) which encode conserved structural proteins (spike protein [S], envelop protein [E], membrane protein [M] and nucleocapsid protein [N]), and several accessory proteins (3a, 6, 7a, 7b, 8 and 10) (39). During its life cycle, SARS-CoV-2 extensively interacts with host factors in order to facilitate cell entry, transcription of viral RNA and translation of subgenomic mRNAs, virion maturation and evasion of the host’s immune response (90; 11; 20). Mechanisms of virus-host interaction are multifaceted and include protein-protein interactions (PPIs), binding of viral proteins to the host transcriptome (96), RNA-RNA interactions and binding of host proteins to viral RNAs. Studies on SARS-CoV-2 infected cells to date have predominantly focused on the entry of SARS-CoV-2 into human epithelial cells, which involves the interaction of the viral spike protein S with the human ACE2 receptor (39). Other studies characterized changes in the host cell transcriptome and proteome upon infection and identified host factors essential for viral replication via CRISPR screenings (78; 25; 92). Lastly, mapping of protein-protein interactions (PPIs) between viral and host proteins has revealed cellular pathways important for SARS-CoV-2 infection. For instance, a recent study identified close to 300 host-virus interactions in the context of SARS-CoV-2 (25). However, these studies have been of limited impact with respect to revealing how the viral RNA is regulated during infection.

RNA viruses hijack key cellular host pathways by interfering with the activity of master regulatory proteins, including RNA binding proteins (RBPs) (29). RBPs are a family of proteins that bind to RNA molecules and control several aspects of cellular RNA metabolism, including splicing, stability, export and translation initiation. In most cases, RNA targets of an RBP share at least one common local sequence or structural feature – a so-called motif - which facilitates the recognition of the RNA by the protein. Host cell RBPs have previously been reported to interact with viral RNA elements and influence several steps of the viral life cycle, such as recruitment of viral RNA to the membrane and synthesis of subgenomic viral RNAs (47; 48; 59; 21). Indeed, in a recent proteome-wide study, 342 RBPs were identified to be annotated with gene ontology (GO) terms related to viruses, infection or immunity with a further 130 RBPs being linked to viruses in literature (21). Examples include the Dengue virus Manokaran et al. (56), the Murine Norovirus (MNV) (88) and Sindbis virus (SINV), where it has been shown that RBPs stimulated by the infection redistribute to viral replication factories and modulate the success of infection (21). The ability of viral RNAs to recruit essential host RBPs could explain permissiveness of certain cell types as well as its range of hosts (48), which is especially relevant for zoonotic viruses such as SARS-CoV-2. In the context of SARS-CoV infection, DEAD-box helicase 1 (DDX1) RBP has been shown to facilitate template read-through and thus replication of genomic viral RNA, while heterogeneous nuclear ribonucleoprotein A1 (hnRNPA1) might regulate viral RNA synthesis (20; 54; 94). Multiple recent studies show that SARS-CoV-2 RNAs extensively interact with both pro-and anti-viral host RBPs during its life cycle (18; 69; 46; 43). Using comprehensive identification of RNA-binding proteins by mass spectrometry (ChIRP-MS), Flynn et al. (18) identified a total of 229 vRNA-bound host factors in human Huh7.5 cells with prominent roles in protecting the host from virus-induced cell death. Schmidt et al. (69) identified 104 vRNA-bound human proteins in the same cell line via RNA antisense purification and quantitative mass spectrometry (RAP-MS), with GO-terms strongly enriched in translation initiation, nonsense-mediated decay and viral transcription. The authors further confirmed the specific location of vRNA binding sites for cellular nucleic acid-binding protein (CNBP) and La-related protein 1 (LARP1) via enhanced cross-linking immunoprecipitation followed by sequencing (eCLIP-seq), which were both associated to restriction of SARS-CoV-2 replication (69). Lee at al. (46) identified 109 vRNA-bound proteins via a modified version of the RAP-MS protocol and linked those RBPs to RNA stability control, mRNA function, and viral process. Further, the authors showed 107 of those host factors are found to interact with vRNA of the seasonal betacoronavirus HCoV-OC43, suggesting that the vRNA interactome is highly conserved. Finally, Labeau et al. (43) used ChIRP-MS to identify 142 host proteins that bind to the SARS-CoV-2 RNA and showed, in contrast to Flynn et al. (18), that siRNA knockdown of most RBPs cellular expression leads to a significant reduction in viral particles, suggesting that the majority of RBPs represent pro-viral factors. Taken together, there is strong evidence that SARS-CoV-2, like other RNA viruses, heavily relies on the presence of a large number of essential RNA-binding host factors. However, the sets of SARS-CoV-2 relevant RBPs from different studies have limited overlap and the outcome depends on the specific cell line utilized in the experiment. Further, most studies lack information of of exact binding sites of human RBPs on viral RNA. A comprehensive large scale analysis of the propensities of different host RBPs to bind to RNA elements across the SARS-CoV-2 genome is currently missing.

Cross-linking and immunoprecipitation (IP) followed by sequencing (CLIP-seq) assays (26), including PAR-CLIP and eCLIP protocols, are the most widely used methods to measure RBP-RNA interactions *in vivo* at high nucleotide resolution and are able to provide sets of functional elements that are directly bound by an RBP of interest (85). While CLIP-seq experiments allow for precise identification of host factor interaction with viral RNAs, the high cost of profiling interactions across a large number of RBPs becomes prohibitive at larger scales, as dedicated pull-down and sequencing has to be performed for each RBP individually. Therefore, such datasets have been generated only for a small number of proteins on SARS-CoV-2 (69). Further, in order to keep up with the continuous emergence of novel SARS-CoV-2 variants, CLIP-seq experiments would need to be repeated for the genome of each viral strain in order to account for (or to identify) gain-or loss-of-binding variants. Recent advances in machine-and deep-learning have enabled a cheaper but powerful alternative by computationally modeling the binding preference of RBPs using information from existing CLIP-seq datasets, such as those generated as part of the ENCODE project (86).

In this study, we train and optimize two recent Convolutional Neural Network (CNN) based methods, Pysster (5) and DeepRiPe (23), on hundreds of human eCLIP and PAR-CLIP datasets and use trained models to predict RBP binding on viral sequences. By that we provide, to our knowledge, the first comprehensive single-nucleotide resolution *in silico* map of viral RNA - host RBP interaction for SARS-CoV-2 as well as 6 other human coronaviruses and identify sequence variants which significantly alter RBP-RNA interaction across 11 different SARS-CoV-2 variants-of-concern. We recapitulate human RBPs which are predicted or experimentally determined to binding to SARS-CoV-2 by previous studies and identify novel host RBP candidates with no previously reported binding to SARS-CoV-2. We integrate knowledge of these proteins across other pathogens and highlight RBPs with clinical relevance, by annotating those that were found among SARS-CoV-2-associated genes from Genome Wide Association Studies (GWAS) (64), CRISPR studies (24; 30; 70; 91), physical binding experiments (18; 69; 89), or patient OMICS data from blood serum and plasma (10; 12; 13; 22; 57; 63; 71; 95). Finally, we perform extensive *in silico* single-nucleotide perturbations across the SARS-CoV-2 genome to identify variants that would lead to gain and/or disruption of RBP binding sites and thus may alter viral fitness.

## 2 Results

The overall workflow of our approach is summarized in Figure 1, from model training, to the *in silico* mapping of the SARS-CoV-2 RBP-RNA interactome and downstream analysis. We first obtained binding site information of publicly available eCLIP experiments of 150 RBPs from the ENCODE (86) database and pre-processed them to obtain a set of high-quality sites of protein-RNA interaction. For each RBP, a convolutional neural network (CNN) classifier to predict the likelihood of RBP-binding to an arbitrary input RNA sequence was trained using the *pysster* (5) framework, resulting in 150 pysster models (Figure 1a). For RBPs not contained in the ENCODE dataset, we included DeepRiPe (23) models pre-trained on 59 PAR-CLIP datasets Next, we performed extensive model performance evaluation on custom trained pysster models and removed poorly performing models from downstream analysis. Using high-quality models, we predicted the likelihood of each RBP binding to individual nucleotides in the SARS-CoV-2 genome using a sliding-window scanning approach (Figure 1b, Methods 3.6). Single-nucleotide binding predictions were further annotated with empirical p-values to correct for false positive hits; and consecutive high-scoring and significant position were aggregated into larger binding-site regions. We thus constructed a comprehensive *in silico* binding map of human RBPs on the SARS-CoV-2 genome and clustered RBP binding sites across different viral genomic regions to unravel potential regulatory patterns (Figure 1b). Exploiting the capability of CNNs to learn complex sequence patterns, we additionally validated our approach by identifying known binding motifs at predicted RBP binding sites. Finally, we utilize our models to score the impact of sequence variant identified in 11 viral strains and identified conserved and novel binding sites across 6 other coronaviruses, including SARS-CoV-1 and MERS (Figure 1c).

**Figure 1:**
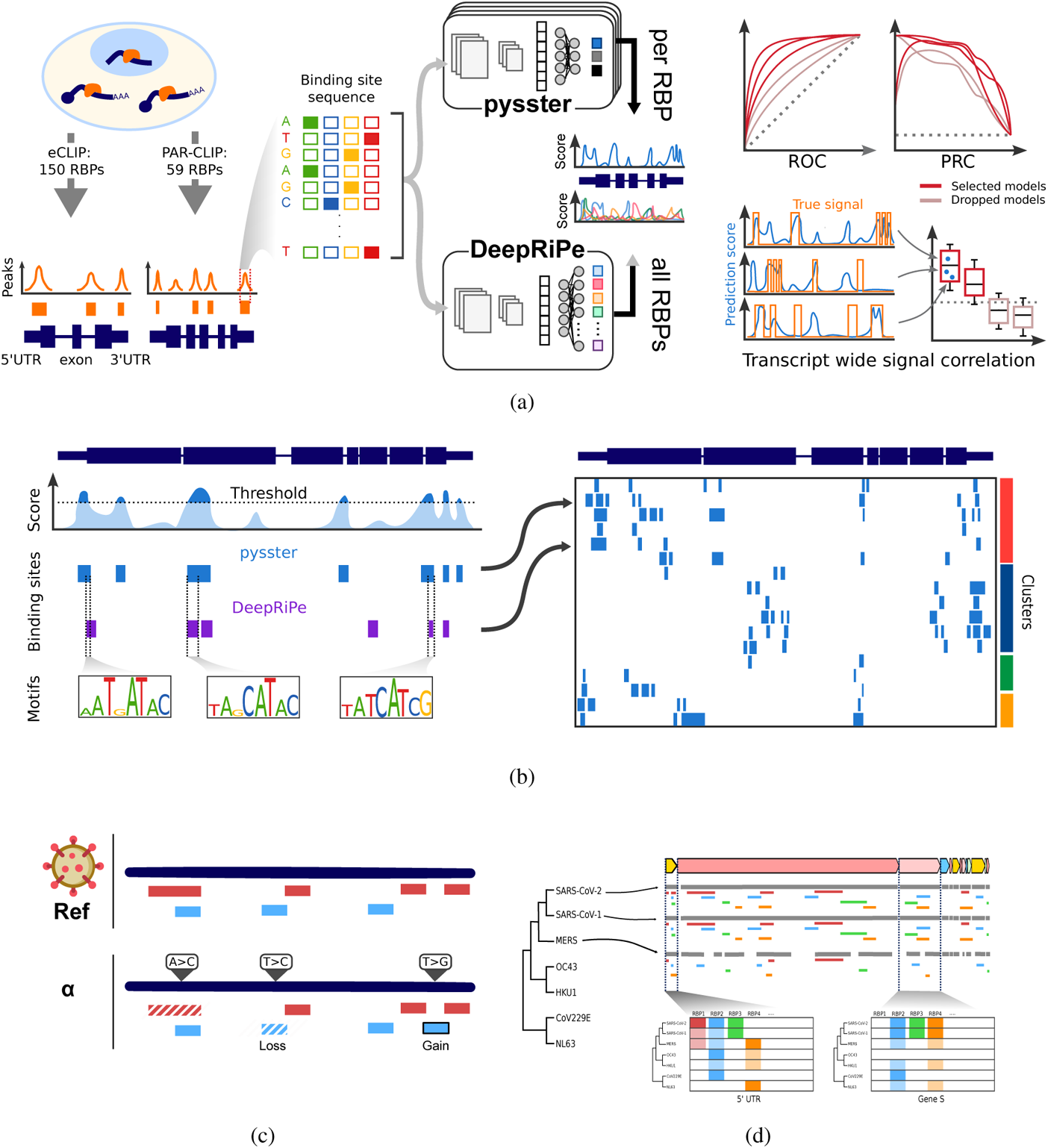
Pipeline of the computational mapping of the human - SARS-CoV-2 protein-RNA interactome. **a**. (Left panel) Interactions between RNA-binding proteins (RBPs) and transcripts can be experimentally measured through eCLIP and PAR-CLIP protocols, enabling the quantification of locally accumulated reads, and the calling of peaks. Such peaks were obtained for 150 RBPs from eCLIP data (86), and for 59 RBPs from PAR-CLIP data (61). (Middle panel) Sequences from these peaks were used to train two deep learning models, composed of convolutional neural networks enabling the detection of complex sequence motifs. These models can then be applied to predict for a given sequence its potential for binding by a RBP. The pysster models are trained separately for each RBP, while DeepRIPE is trained in a multi-task fashion and simultaneously for all input RBPs. (Right panel) A selection of high-performance models was established through evaluation of performance of the models, from overall performance metrics to in-practice, sequence-wide evaluation. **b**. All retained models were applied to scan the entire genome of SARS-CoV-2, and binding sites were predicted from consecutive, high-prediction scores positions. Sequence motifs underlying RBP binding sites were also identified by interrogating both CNNs via Integrated Gradients. Predictions were compiled in the first *in silico* map of host-protein - viral RNA interactome for SARS-CoV-2. **c** The prediction models were applied to evaluate the impact of variants of concerns, **d** as well as to evaluate the evolutionary trajectory of affinity of host RBPs to other coronaviruses’ genomes.

### 2.1 Accurate model predictions in human and viral sequences

The trained pysster models showed a robust area-under-precision-recall-curve (auPRC) performance (Methods 3.7.1), with a median auPRC of 0.6 across all 150 trained models (Figure 2a). As models were used for scanning of the full-length viral genome (rather than classification of standalone examples), we further evaluate the model performance by computing the correlation of the predicted positive-class probabilities with observed ENCODE peaks on a hold-out set of human transcripts (Methods 3.7.2). Nearly all models showed a significant positive correlation, with a mean median Spearman correlation coefficient (SCC) across transcripts of 0.149 and a maximum median SCC of 0.38 (Figure 2b), indicating that the trained models are well-suited for the task of scanning across the viral genome. Exemplary prediction tracks for two held-out human transcripts using pysster models of QKI and TARDBP are shown in Figure 2c. In general, we observe that models which perform well with respect to the auPRC score tend to perform well in the context of RNA sequence scanning (Figure 2d). To ensure that downstream analyses are based on a high-quality set of binding site predictions, models with a median SCC of less than 0.1 or an auPRC of less than 0.6 were discarded (Methods 3.7.2). A total of 63 high-quality pysster models were thus kept for predicting on the SARS-CoV-2 genome. For DeepRiPe, we relied on the results from (23) and retained only those models where informative sequence motifs were learned during training, leaving a total of 33 RBP models for predicting on the SARS-CoV-2 genome. Of those, we selected only models for RBPs not contained in the ENCODE database, leading to the addition of 24 high-quality DeepRiPe models. To confirm that pysster models trained on CLIP-seq data from human cell lines are suitable for cross-species binding-site inference in SARS-CoV-2, we validated our approach for RBPs with available CLIP-seq experiments from SARS-CoV-2 infected human cell lines. To this end, we obtained eCLIP datasets for CNBP and LARP1 on both human and SARS-CoV-2 transcripts from Schmidt et al. (69) and processed binding sites as described in Section 3.1. After generating training samples on CNBP and LARP1 binding sites within human transcripts (Methods 3.2), we trained pysster models for both RBPs. We then performed prediction along the SARS-CoV-2 RNA sequence and compared the resulting prediction scores with observed binding sites as well as the raw eCLIP signal (Figure 2e, 2f). Predictions from pysster models trained on human binding sites showed a strong correlation with the raw eCLIP signal (SCC = 0.332, p-value < 1e-16 for CNBP and SCC = 0.133, p-value = 7.96e-12 for LARP1), and accumulation of high-scoring positions at the location of called binding sites from the eCLIP experiment (Figure 2f). Further, we observed significantly higher prediction scores for in-binding-site nucleotides versus outside-binding-site nucleotides for both RBPs (Figure 2f; t-test, p-value < 1e-16 for CNBP; p-value = 2.44e-6 for LARP1). Taken together, these results strongly support the validity of our approach for cross-species *in silico* prediction of RBP binding sites.

**Figure 2:**
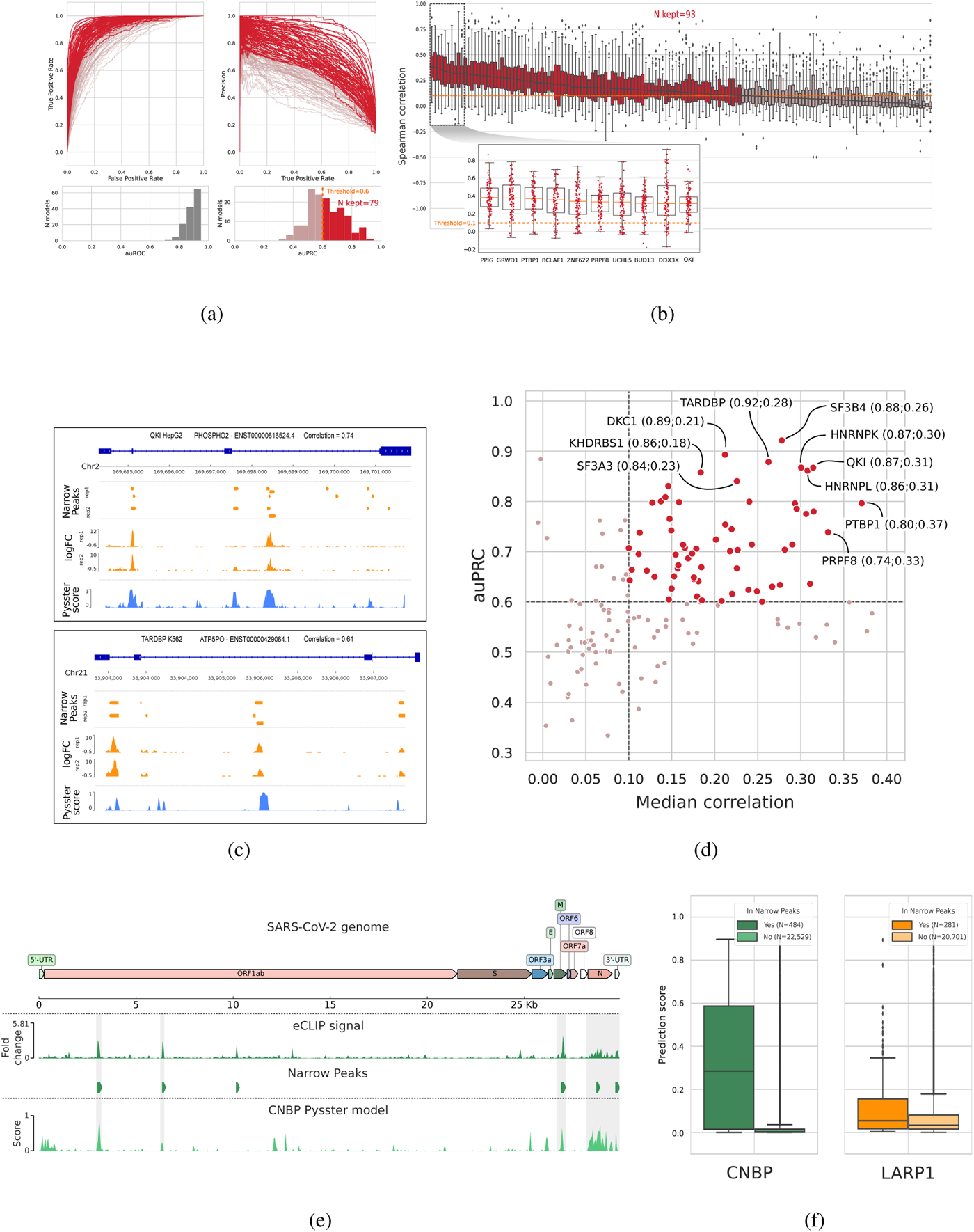
Evaluation of pysster models’ performance and high-quality model selection. **a**. Receiver Operating Curve (ROC) and Precision Recall Curve (PRC) for all 150 pysster models trained from ENCODE eCLIP datasets. A first threshold of 0.6 was set on the area under the PRCs (auPRC), leading to a subset of 79 models passing the threshold. **b**. Boxplots of correlations between eCLIP and prediction scores from 100 left-out transcripts per RBP model. This correlation highlights the performance of models in a realistic context of full-sequence-length scan. A second threshold was thus set on the median correlation coefficient, leading to a subset of 93 models passing the threshold. The 10 models with highest median correlation are displayed in a detailed sub-plot. **c**. Genome-browser view illustrating the comparison between eCLIP signals and model prediction scores over full-length transcripts. Two of the best models are presented, with signal from left-out transcripts with high correlation between eCLIP log-fold-change signals and prediction scores from the pysster models. **d**. Scatterplot of the AUPRC and median correlation values for each model, highlighting the final subset of high-quality models. The top 10 models are labeled. **e**. Comparison of genome-wide eCLIP signal and pysster prediction scores from the CNBP eCLIP datasets generated over the SARS-CoV-2 genome by (69). **f**. Boxplot of pysster prediction scores from position within or without overlap from called narrow peaks, for the CNBP model and the LARP1 model.

### 2.2 A comprehensive *in silico* binding map of human RBPs on SARS-CoV-2

We performed *in silico* binding site calling by identifying consecutive significant and high-scoring positions within the SARS-CoV-2 genome with both pysster and DeepRiPe high-confidence models (Methods 3.9). In the following, we first demonstrate that our model predictions correspond to *bona fide* RBP binding sites on the SARS-CoV-2 genome by performing motif analysis and subsequently build a computational map of SARS-CoV-2-human RBP interactions. We then evaluate the enrichment of different RBPs for different viral genomic regions, as well as their putative regulatory function in the context of SARS-CoV-2 infection.

#### Predicted RBP binding sites coincide with known binding motifs

Figure 3a and 3b each show single-nucleotide resolution prediction scores of the well-known human RBPs RBFOX2 and TARDBP, obtained from pysster models, and MBNL1 and QKI, obtained from DeepRiPe models. Identified binding sites (Methods 3.9) are shown below the prediction score tracks. To identify driving features of RBP binding and to ensure that high-scoring positions represent genuine binding sites rather than model artifacts we performed feature importance analysis (Methods 3.10) in order to assess whether the sequence features underlying the predictions at those sites correspond to the binding site preferences of those proteins reported in literature. Specifically, we centered input windows around predicted binding sites of RBFOX2, TARDBP, MBNL1 and QKI on SARS-CoV-2 to identify individual nucleotides that were most predictive for classifying the input sequence as ‘bound’ (Figure 3a and 3b; bottom track). We observed that feature importance maps around predicted binding sites corresponded to known binding motifs. For instance, we observe the well-known consensus sequence (T)GCATG recognized by the splicing factor RBFOX2 (36) in the corresponding feature importance maps (Figure 3a, left), as well as the TG-repeat motif, corresponding to the sequence preference of TARDBP (28), coinciding with its predicted binding sites (Figure 3a, right). Similarly, DeepRiPe attribution maps with respect to binding sites of QKI show the canonical binding motif TACTAA(C) (82) (Figure 3b, left). Lastly, the attribution maps computed at each binding site of the splicing factor MBNL1 all harbour occurrences of the characteristic YGCY motif (45) (Figure 3b, right).

**Figure 3:**
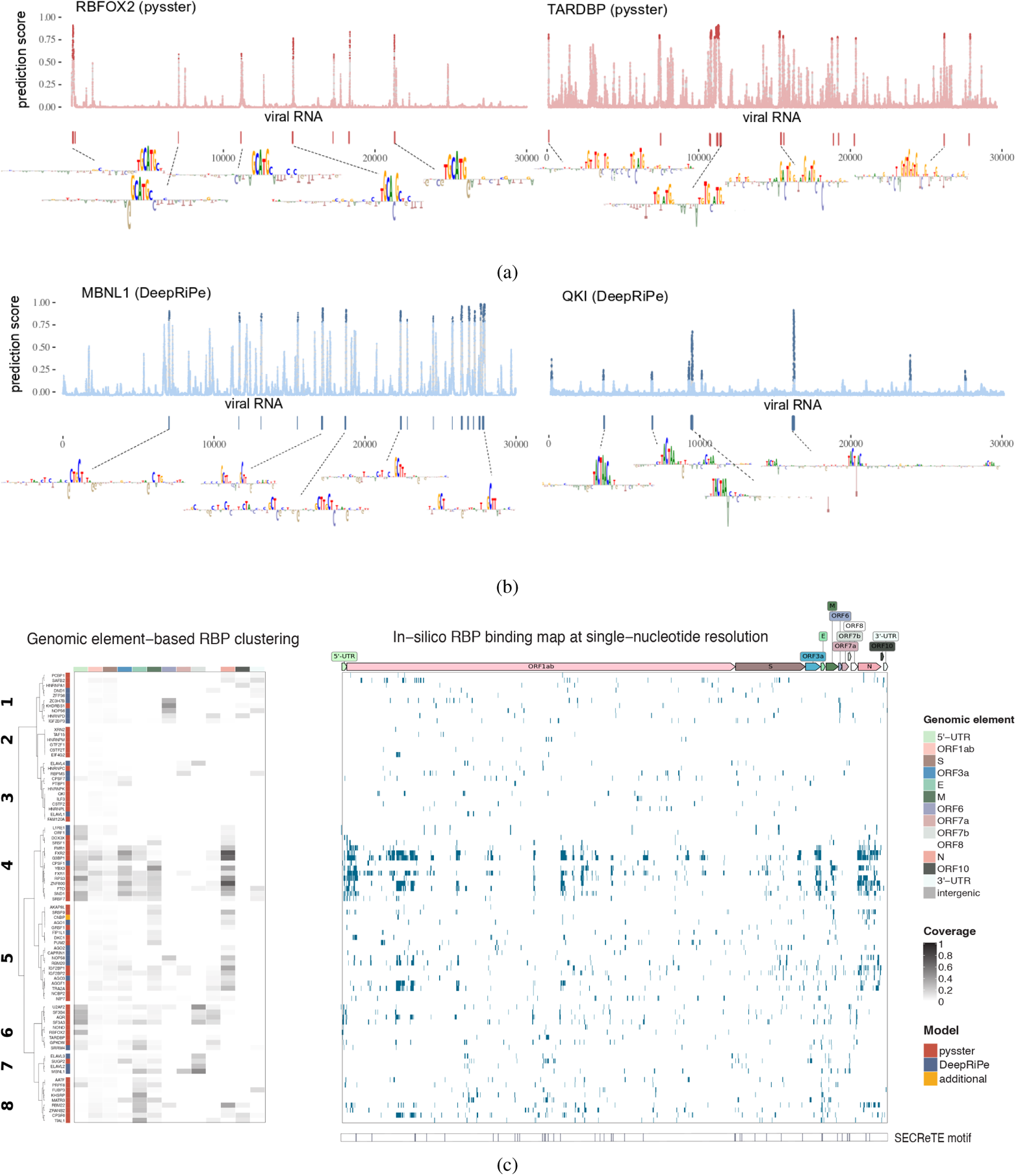
Computational map of RBP binding on SARS-CoV-2. **a** Single-nucleotide probability score for RBFOX2 (left) and TARDBP (right) RBP binding as computed by the corresponding pysster models across the whole SARS-CoV-2 genome. The higher the score, the higher the likelihood of a binding event at that position. Points highlighted in strong color correspond to significant predictions, i.e. with bound probability significantly higher than random (empirical p-value *<* 0.01, see Methods). Wider binding sites, encompassing more than one significant position are shown as vertical bars underneath each prediction profile, together with their corresponding binding motifs as extracted by means of attribution maps (see Methods). **b** Single-nucleotide probability score for MBNL1 (left) and QKI (right) RBP binding as computed by the corresponding DeepRiPe models. Significant positions (empirical p-value *<* 0.01) are highlighted in strong color, and computed binding sites together with their corresponding motifs are shown underneath. **c** Clustering of RBPs based on binding site coverage of genomic annotations of SARS-CoV-2 for both pysster and DeepRiPe RBPs (left panel). *In silico* RBP binding map, at single-nucleotide resolution, for both pysster and DeepRiPe RBPs (right panel). SARS-CoV-2 SECReTE motifs from (27) are shown below.

#### Binding site predictions are robust across different datasets and prediction tools

To evaluate the robustness of viral binding site predictions across pysster and DeepRiPe, we compared predictions for a small set of RBPs where both eCLIP data (used to train a pysster models) and PAR-CLIP data (used for the training of DeepRiPe models) were available. Among a total of 20 overlapping RBPs, 12 were contained in the sets of high-quality models for pysster and DeepRiPe selected in 2.1, namely TARDBP, CSTF2, IGF2BP1, PUM2, CSTF2T, QKI, IGF2BP2, IGF2BP3, CPSF6 FXR1, FXR2 and EWSR1. For each of the 12 RBPs, we then computed the Spearman correlation between the pysster and DeepRiPe prediction scores across single-nucleotide positions on the viral genome. We observed a signal correlation higher than 0.1 for 8 out of the 12 RBPs, with a Spearman correlation coefficient ranging from a maximum of 0.64 (TARDBP) to a minimum of 0.15 (CPSF6) (Supplementary Table 1). In general, we observed a higher overlap between pysster and DeepRiPE binding site predictions for RBPs harbouring well-defined RNA sequence motifs, such as QKI, TARDBP, PUM2, CSTF2, and to a less extent, FXR1/2 and IGF2BP1/2/3. In addition, feature attributions maps at overlapping binding sites of pysster and DeepRiPe with respect to QKI and TARDBP (Supplementary Figure 1), highlight the presence of the known binding motifs for these two RBPs.

#### Binding preferences and clusters of human RBP predicted sites on the SARS-CoV-2 genome

Given the strong evidence that our predictions reflect true likelihoods of viral sequence regions being bound by human RBPs, we set out to build a full *in silico* SARS-CoV-2 / human RBP binding map, using the set of 88 high confidence models from both pysster and DeepRiPe (Section 2.1). Note that we included the CNBP model from Section 2.1, as it satisfied our performance constrains. Further, for the 12 shared RBPs between pysster and DeepRiPe, only pysster predictions were considered for downstream analysis, given the high agreement between both models. Figure 3c (right) depicts the binding profiles of 84 (out of 88) human RBPs which harbor at least one binding site on the SARS-CoV-2 sequence. We clustered RBPs into eight classes based on their relative binding site coverage across different genomic regions of the SARS-CoV-2 genome (Figure 3c, left). We observe that some clusters of proteins exhibit sparse binding signal across the SARS-CoV-2 genome (such as clusters 2 and 3), while other clusters contain RBPs which are predicted to bind extensively across the whole SARS-CoV-2 genome (cluster 4). Interestingly, some clusters harbour RBPs shown to preferentially bind specific genomic elements (cluster 1 and cluster 5-8, Figure 3c, left). We observe overall extensive RBP binding coverage mostly at 5’ UTRs and genomic regions coding for E, M and N structural proteins, and less coverage at the spike S gene, as well as the viral 3’ UTR. To some extent, clustering of predicted binding sites groups together RBPs with similar functions in RNA processing and viral regulation, as well similar RNA recognition mechanisms. Cluster 4 corresponds to a group of well-known regulators of RNA processing, which extensively bind the viral 5’ UTR, as well as the ORF1ab and subgenomic RNAs. This includes proteins from the FXR family (FXR1, FXR2 and FMR1), which recognize RNA using the K Homology (KH) domain, and control RNA stability, translation and RNA localization (85). Other RNA translational regulators in the same cluster include the DDX3X helicase, which was recently identified as host target against SARS-CoV-2 infection (9), and the 40S ribosomal protein S3 (RPS3), which also binding RNAs through the KH domain. Other proteins in this cluster with well-known roles in regulation of viral infections are SND1, the splicing regulators (SR) SRSF1 and SRSF2, shown to be implicated in increasing translation efficiency in the context of HIV infection (55), the RNA demethylase factor FTO, known to regulate viral infections and HIV-1 protein expression (83), in addition to the aforementioned G3BP1 and DDX3X involved in innate immunity (8). Cluster 1 predominantly harbors RBPs with binding preference for the viral 3’ UTR, including regulators of RNA stability and proteins involved in 3’ end formation and/or regulation of translation. Among those RBPs, the poly (I:C) binding protein KHDRBS1 has been identified to have pro-viral activity in SFV infection (65), while the multifunctional RBP PCBP1, along with hnRNPRs has been shown to be implicated in translational control of many viruses, including poliovirus, human pailloma virus and Hepatitis A virus. Cluster 6 is comprised of RBPs which preferentially bind to the 5’ UTR of SARS-CoV-2. Interestingly, these proteins (AQR, GPKOW, SF3A3, SF3B4 and A2AF2) are known to be functionally involved in splicing and harbour a RNA recognition motif (RRM) (85). We find that cluster 6 also harbors NONO, a member of the paraspeckle complex, which has previously been associated with antiviral immune response and which is part of the RBP interactome in SINV infected cells (21), as well as TARDBP, a protein that localizes to P-bodies and stress granules and was shown to bind to the 5’ UTR of SARS-CoV-2 in a recent study (60). Cluster 5 includes a large class of RBPs with diverse functions, including splicing (SRSF9), post-transcriptional repression (PUM2 and CAPRIN1), snoRNA binding (NOP58 and NIP7) and miRNA-mediated silencing (AGO1-3). These proteins were predicted to preferentially bind to the N and M genomic regions, while being depleted in the viral UTRs.

Lastly, binding of RBPs in cluster 7 and 8 is mostly concentrated in ORF7b as well as E and M protein regions, respectively. Besides the splicing regulators MBNL1 and SUGP2, cluster 7 contains the ELAVL2 and ELAVL3 RBPs involved in regulation of RNA stability (38). Previous studies have suggested that ELAVL human proteins might be affected during infections by the viral RNA that acts as a competitor to tritate them away from their cellular mRNA targets (66). While most RBPs in cluster 8 were not found to be functionally related in literature, RBPs KHSRP and MATR3 have been shown to act as restriction factors in SINV infection (65)

#### Predicted RBP binding sites overlap with SECReTE motifs

Haimovich et al. (27) recently identified the presence of a unique *cis*-acting RNA element, termed “SECReTE” motif, which consists of 10 or more consecutive triplet repeats, with a C or a U present at every third base, on the sequences of both (-) and (+)ssRNA viruses. In context of SARS-CoV-2, a total of 40 SECReTE motifs have been identified in the viral genome, with a total length of ~1.3 kilobase. This motif has been suggested to be important for efficient translation and secretion of membrane or ER-associated secreted viral proteins, as well as for viral replication centers (VRCs) formation. To investigate whether predicted binding sites identified in 2.2 coincide with SECReTE motifs, we obtained exact locations of all SARS-CoV-2 SECReTE motifs from (27), and subsequently intersected them with predicted RBP binding sites of all 84 high-quality models containing at least one binding site in SARS-CoV-2. We observed that a total of 61 RBPs (out of 84) have binding sites overlapping with SECReTE motifs. Further, 30 RBPs with at least 10% of their binding sites overlapping with SECReTE motifs were identified and are termed ‘SECReTE-associated RBPs’ subsequently. We find that SECReTE-associated RBPs are predominantly found in some clusters of Figure 3c (cluster 3 and 6-8), while showing an apparent depletion in others (cluster 1-2, Figure 3c). For instance, 5 (out of 9) SECReTE-associated RBPs (SF3B4, U2AF2, GPKOW, TARDBP and NONO) are found in cluster 6, with TARDBP and NONO being functinally associated to viral regulation (85; 65). Cluster 3 contains 5 (out of 12) SECReTE-associated RBPs, namely CSTF2, ELAVL4, HNRNPC, PTBP1 and QKI, each associated with multiple RNA functional processes, including RNA stability, 3’-end formation, splicing and translation (85). Cluster 8 harbors 4 (out of 9) SECReTE-associated RBPs (FUBP3, KHSRP, MATR3 and CPSF6), 3 of which (FUBP3, KHSRP, MATR3) have 25% or more of their binding sites overlapping with SECReTE motifs. KHSRP is an essential RBP involved in RNA localization, RNA stability and translation, while METR3 is a regulator of RNA stability. Interestingly, most of these factors have been previously associated to viral RNA regulation (85). Lastly, all 4 RBPs in cluster 7 (ELAVL2, ELAVL3, SUGP2 and MBNL1) appear to be strongly associated with SECReTE motifs, as more than 25% of their respective binding sites are overlapping genomic regions harbouring SECReTE motifs.

### 2.3 SARS-CoV-2 variants of concern show gain- and loss-of-binding events

Multiple waves of SARS-CoV-2 infections have spread across the globe, some of which resulted in the emergence of specific lineages of viral variants. The systematic sequencing of thousands of samples from infected patients enabled the description and categorization of the detected viral sequences, identifying numerous mutations in their sequence when compared to the initial SARS-CoV-2 reference genome. Some of the thus described strains have been experimentally characterized as more efficient than others, explaining in part their successful spread at local or global geographic scales (32; 84; 33). These strains have been defined by the World Health Organization as variants of concern, with “evidence for increased transmissibility, virulence, and/or decreased diagnostic, therapeutic, or vaccine efficacy” (67). Specific subsets of mutations have been associated with each variant of concern, when mutations were represented in a majority of sequenced samples of their lineage. Notably, a special focus has been given with regards to the impact of mutations occurring within the spike-encoding gene (50), owing its importance in the initial steps of viral infection and its potential for vaccine neutralization (31). However, due to a lack of appropriate methods, the impact of these mutations at the regulatory level, such as their impact on protein-RNA interactions, has so far been largely ignored.

To fill this gap, we systematically investigated the impact of observed mutations in viral variants of concern on the predicted binding of RBPs, in order to uncover potential viral hijacking of host proteins directly at the RNA level.

#### A catalog of high-impacting variants across 11 viral strains

We compiled a total of 290 mutations (193 unique mutations, 37 shared across strains) across 11 variants of concern, including alpha, delta, and omicron strains (Methods 3.11). For each variant and RBP, we evaluated the impact of the variant in terms of gain- or loss-of-binding by comparing the predicted binding probability of the reference and alternative allele (Methods 3.11.) Using pysster and DeepRiPe models across 87 RBPs, we obtained a total of 25,230 impact scores, one for each pair of variant and RBP. Notably, three variants (3,037C>T, 14,408C>T, and 23,403A>G) are consistently found across all viral strains, and their highest absolute delta-scores were respectively associated to FTO (avg. decrease from 0.474 to 0.356), AQR (avg. decrease from 0.191 to 0.036), and NONO (avg. increase from 0.086 to 0.340). In order to prioritize pairs of variants and RBPs that show a gain- or loss-of-binding, we select a sub-set of pairs for which either the reference *or* alternative allele pass our binding thresholds (Methods 3.9). Note that this filter applies a XOR operation, i.e. we are interested in events that lead to either gain- or loss-of-binding (GOB, LOB). Overall, a total of 315 GOB or LOB events passed the above filter and are depicted in Figure 4a. The majority of variants introduced small delta in prediction scores, with less than 20% (61) of absolute delta-scores above 0.233 (Figure 4a). As shown in the Supplementary Figure 2a, the top 20% highest-impact variants from Figure 4a accumulate in different genomic annotations over the SARS-CoV-2 genome. Interestingly, among the RBPs impacted by these mutations, we find that some strains present multiple high-delta-score mutations for SRSF7 (strains delta, kappa) and YBX3 (strain lambda), as well as L1RE1, RBPMS, SND1, ZRANB2 (strain omicron) (Supplementary Figure 2b). Additionally, the omicron strain harbors a particularly large number of variants predicted to impact binding of ORF1 protein (from LINE-1 retrotransposable element).

**Figure 4:**
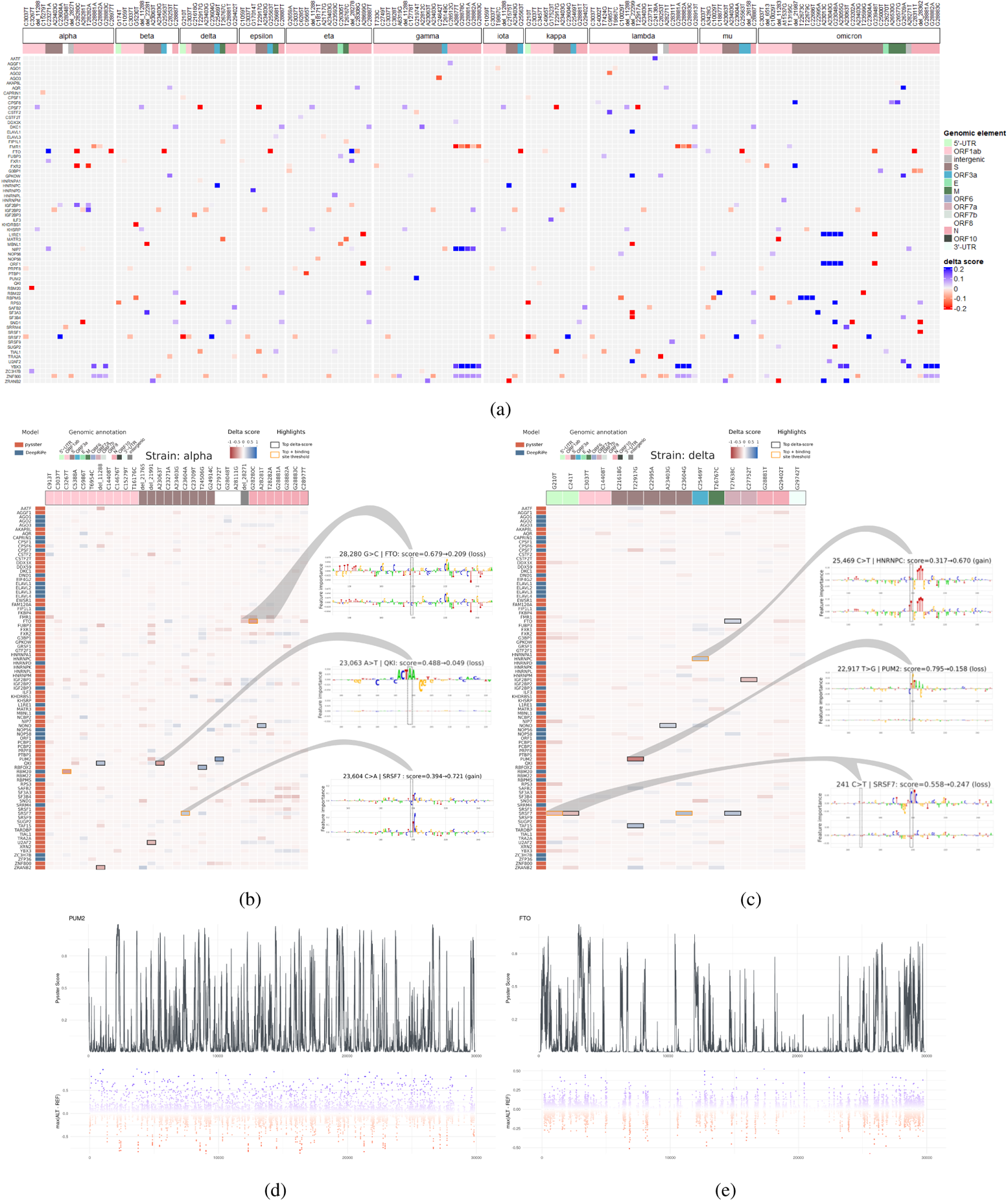
Impact of variants from SARS-CoV-2 strains on predicted binding sites. **a**. Joint heatmap of delta scores from the 290 identified variants in the different SARS-CoV-2 strains. Delta-scores represent the difference in prediction score of a prediction model between alternative and reference sequences centered on each variant. Only the 315 impacts labeled as change-of-binding are colored (see 3.11). Delta score color scale is capped so as to show low delta score impacts. RBPs and mutations without any such impact across strains are dropped from the heatmap. **b**. Complete heatmap of delta scores from 31 variants associated to the alpha viral variant. The top 10 with highest absolute delta scores are lined out, with yellow color indicating the ones labeled as change-of-binding. Some sites are further investigated through integrated gradients, comparing the sequence motifs identified by the prediction models against known motifs from mCrossBase (16). **c**. Complete heatmap of delta scores from 16 variants associated to the delta viral variant. **d,e**. Results from the *in silico* mutagenesis over the SARS-CoV-2 genome. Nucleotides across the viral genome were perturbed towards the three alternative bases, generating a reference distribution of possible delta-scores, notably highlighting positions with highest impacts. Here, **d)** and **e)** display the position-wise reference score (top) and delta score (bottom) for PUM2 and FTO, respectively.

#### Systematic point-wise *in silico* mutagenesis reveals hypothetical high-impact variants

New viral strains are continuously emerging, some of which are characterized by a faster spread due to newly acquired sequence variants, highlighting the importance of a continuous monitoring of viral variants which may result in a selective advantage on the protein or RNA regulatory level. To anticipate and quantify the impact of potentially unobserved variants, we perform a systematic *in silico* mutagenesis by generating all possible point mutations across the SARS-CoV-2 genome and score each hypothetical mutation with respect to its impact on RBP binding. Figure 4d and 4e show exemplary *in silico* mutation tracks for PUM2 and FTO, respectively, with observed reference prediction scores depicted at the top and the impact of gain- and loss-of-binding variants shown at the bottom. Note that for visualization purpose, only the delta score of the alternative allele with the highest impact is shown for each position and RBP. Supplementary Figure 3 shows an impact catalogue of 29, 903 *×* 63 single-nucleotide variants across all SARS-CoV-2 genome positions and 63 pysster models. The complete set of hypothetical variants together with their impact scores is available at https://sc2rbpmap.helmholtz-muenchen.de/.

#### High-impact sequence variants disrupt known RBP-binding motifs

As *in vivo* RBP-binding is usually driven via the recognition of short sequence motifs, we investigated whether high-impact variants cause gain or disruption of known binding motifs. To this end, we gathered from each strain the top 10 variants with highest absolute delta-scores, as illustrated in Figure 4b and 4c for strains alpha and delta, respectively. This represented a total of 69 unique mutation-RBP pairs, 19 of which were found in more than one strain. As expected, the majority (54 / 69) of their delta-scores is found to be in the top 1% of the distributions from the *in silico* mutagenesis. We then computed feature attribution scores (Methods 3.10), centered at the position of each high-impact variant. Feature attribution maps for the subset of candidate high-impact variants of the alpha and delta strain are shown in Figure 4b and 4c, respectively. Indeed, we observe that variants with high negative delta score tend to disrupt known binding motifs of human RBPs. For instance, transition T>G at position 22,917, as seen in the delta strain (Figure 4c) (as well as in top mutations from epsilon and kappa strains) decreases the prediction score for PUM2 from 0.795 to 0.158, with only 0.0015% in silico variants showing a lower delta-score. As is clearly visible from the feature attribution analysis (Figure 4c; middle-right), the variant disrupts the well-known PUM2 binding motif TGTATAT. In a similar manner, transversion A>T at position 23,063 from the alpha strain (Figure 4b; also found in top mutations from beta, gamma, and mu strains) decreases the prediction score for QKI from 0.488 to 0.049, with 0.006% *in silico* mutations show a low delta-score. Here, the feature attribution profiles clearly highlight how the known QKI binding motif ACTAA was detected by the model in the reference sequence, and how the mutation leads to a loss of this motif. Lastly, the transversion G>C at position 28,280 in the alpha strain (Figure 4b) decreases the prediction score for FTO binding from 0.679 to 0.209, and only 6 (0.00007%) *in silico* mutations show a delta-score lower than the one observed (Figure 4d). Although no clear motif is found within the window, the heights of the nucleotides at the position of the mutation are reduced compared to the reference sequence, reflecting the decreased prediction score.

#### High-impact gain- and loss-of-binding events across viral strains

Among the above set of top 10 highest impact variants per viral strain, we select those that conform to strict gain-or loss-of-binding (Methods 3.11). We identify a total of 23 (out of 69) change of binding events across 17 variants and 13 RBPs (Table 1). The first example corresponds to a transversion G>T at position 210 in the 5’UTR from the delta and kappa strains, predicted to induce a loss-of-binding for SRSF7, which we had confirmed from the loss of binding motif (delta strain heatmap, see Figure 4c). Further, from the ORF1ab gene, two examples of a loss of binding for RBM20 by the C>T transition at position 3,267 (strain alpha), and a gain of binding of RBM22 from a C>T transition at position 18,877 (strain mu). From the S gene, a gain of binding is reported for HNRNPC, induced by a C>T transition at position 21,575 (strain iota), in addition to another gain of binding reported for SF3A3, from a C>A transversion at position 22,995 (strain omicron). Two mutations occurring in the ORF3a gene are passing our filters for two RBP impacts: the transition C>T at position 25,469 induces a gain of binding for HNRNPC in delta and kappa strains, while the G>T transversion at position 25,563 induces a loss of binding for FTO in strains beta, epsilon, iota and mu. Finally, in the N gene, we report three mutations, two of them impacting FTO binding (one gain in the eta strain, from a deletion at position 28,278; one loss in the alpha strain, from a G>C transversion at position 28,280), and a loss of binding of ORF1 protein (from LINE-1 retrotransposable element) in the eta strain, from a A>G transversion at position 28,699.

**Table 1:** Subset of high delta score mutations passing binding sites thresholds

|  | RBP | Variant | Strain | Genomic element | REF score | ALT score | delta score | Impact |
| --- | --- | --- | --- | --- | --- | --- | --- | --- |
| 0 | SRSF7 | G210T | delta, kappa | 5' UTR | 0.768 | 0.457 | -0.311 | loss |
| 1 | RBM20 | C3267T | alpha | ORF1ab | 0.813 | 0.336 | -0.477 | loss |
| 2 | RBM22 | C18877T | mu | ORF1ab | 0.338 | 0.614 | 0.276 | gain |
| 3 | HNRNPC | C21575T | iota | S | 0.374 | 0.840 | 0.467 | gain |
| 4 | MBNL1 | del_22281 | beta | S | 0.800 | 0.006 | -0.795 | loss |
| 5 | ELAVL1 | del_22299 | lambda | S | 0.070 | 0.632 | 0.562 | gain |
| 6 | SF3B4 | del_22299 | lambda | S | 0.871 | 0.128 | -0.744 | loss |
| 7 | SF3A3 | del_22299 | lambda | S | 0.860 | 0.273 | -0.587 | loss |
| 8 | U2AF2 | del_22299 | lambda | S | 0.543 | 0.980 | 0.438 | gain |
| 9 | GPKOW | del_22299 | lambda | S | 0.297 | 0.841 | 0.544 | gain |
| 10 | MBNL1 | del_22299 | lambda | S | 0.803 | 0.398 | -0.405 | loss |
| 11 | SF3A3 | C22995A | omicron | S | 0.081 | 0.808 | 0.726 | gain |
| 12 | ORF1 | A23013C | omicron | S | 0.014 | 0.621 | 0.608 | gain |
| 13 | ORF1 | A23040G | omicron | S | 0.006 | 0.673 | 0.666 | gain |
| 14 | ORF1 | G23048A | omicron | S | 0.006 | 0.606 | 0.600 | gain |
| 15 | SND1 | G23048A | omicron | S | 0.187 | 0.791 | 0.604 | gain |
| 16 | SRSF7 | C23604A | alpha, mu | S | 0.394 | 0.719 | 0.326 | gain |
| 17 | SRSF7 | C23604G | delta, kappa | S | 0.394 | 0.792 | 0.398 | gain |
| 18 | HNRNPC | C25469T | delta, kappa | ORF3a | 0.317 | 0.670 | 0.352 | gain |
| 19 | FTO | G25563T | beta, epsilon, iota, mu | ORF3a | 0.633 | 0.080 | -0.552 | loss |
| 20 | FTO | del_28278 | eta | N | 0.335 | 0.683 | 0.348 | gain |
| 21 | FTO | G28280C | alpha | N | 0.679 | 0.209 | -0.470 | loss |
| 22 | ORF1 | A28699G | eta | N | 0.597 | 0.141 | -0.456 | loss |

#### Individual variants impact binding of several RBPs

Among variants that surpass binding-sites thresholds and lead to either gain- or loss-of-binding (Methods 3.11), several variants impact RBP binding of multiple RBPs simultaneously. For instance, a deletion at position 22,299 (S gene) identified in the lambda strain, is predicted to induce a gain of binding for ELAVL1, U2AF2, and GPKOW, while inducing a loss of binding for SF3B4, SF3A3, and MBNL1. Interestingly, all these factors are associated with splicing. Notably, the MBNL1 loss is also detected in the beta strain, through a deletion happening in a close-by location (at position 22,281, S gene), suggesting those two mutations may have been retained due to beneficial induction of similar changes in binding patterns. Another mutation which impacts multiple RBPs is the transition G>A at position 23,048 (S gene) from the omicron strain, predicted to induce binding of the ORF1 protein from LINE-1 retrotransposable element, as well as of SND1. Comparably to the MBNL1 impact, two close-by mutations from omicron were associated with a gain of ORF1 binding (transversion A>C at position 23,013, and transition A>G at position 23,040), further suggesting joint impact of these mutations on ORF1p binding. The last case of mutations with impact on multiple RBPs concerns a set of 2 mutations: C>A transversion and C>G transversion at position 23,604, in the S gene. The first is found in alpha and mu strains, while the second is found in the delta and kappa strains. Both mutations are predicted to induce a gain of SRSF7 binding, which is visualized for the alpha strain on Figure 4b through feature attribution maps.

### 2.4 RBP-binding across human coronaviruses

While evaluation of impact for reported variants enables the monitoring of potentially functional changes in the SARS-CoV-2 genome, evaluating changes in binding sites at longer evolutionary time scale might highlight more fundamental properties of the SARS-CoV-2 virus, as compared to other RNA viruses infecting human. We investigated to which extent binding sites of human RBPs are conserved across related human coronaviruses. For this purpose, we obtained genomes and genomic annotations of 6 SARS-CoV-2-related human coronaviruses, namely SARS-CoV-1, MERS, HCoV-OC43, HCoV-NL63, HCoV-HKU1, HCoV-229E (Methods 3.13). Binding sites were identified in analogy to SARS-CoV-2 (Methods 3.9) across each viral genome using 87 high-confidence pysster and DeepRiPe models. Figure 5a shows the general binding propensity of RBPs across viral genomes of the 7 coronaviruses. Overall, RBP binding is conserved across coronaviruses, with the highly pathogenic viruses (SARS-CoV-1, SARS-CoV-2 and MERS) showing a highly similar binding pattern. Further, a total of 86 (out of 87) RBPs (except FKBP4) were predicted to harbor a binding site in at least one coronavirus, with only a small variability in the total number of binding RBPs between individual viruses. However, we observe a greater variability of RBP binding within shared genomic regions across coronaviruses, for instance in the 5’ and 3’ untranslated regions (UTRs). Viral UTRs are known to play an important role in both pro- and anti-viral responses and recent evidence suggests that evolution of the 3’ UTR is contributing to increased viral diversity (15). Indeed, the 3’ UTR of SARS-CoV-2 shows a severe truncation when compared to SARS-CoV-1 and MERS. Given that viral UTRs are not under selective pressure with respect to a translated protein, they might be more prone to acquire mutations that modulate regulation through host RBPs. Figure 5b and 5c show RBP binding to the 3’ and 5’ UTRs across selected coronaviruses, respectively. While SARS-CoV-1, SARS-CoV-2 and MERS show conserved binding on the 5’ UTR and cluster closely, a depletion of RBP binding sites is observed in the 3’ UTR of SARS-CoV-2 when compared to SARS-CoV-1 and MERS. To investigate gain-and loss-of-binding in viral UTRs across the severe pathogenic human coronaviruses SARS-CoV-1, SARS-CoV-2 and MERS, we performed multiple sequence alignment of the viral 3’ and 5’ UTRs and compared the predicted binding score profiles across the three viruses (Methods 3.13).

**Figure 5:**
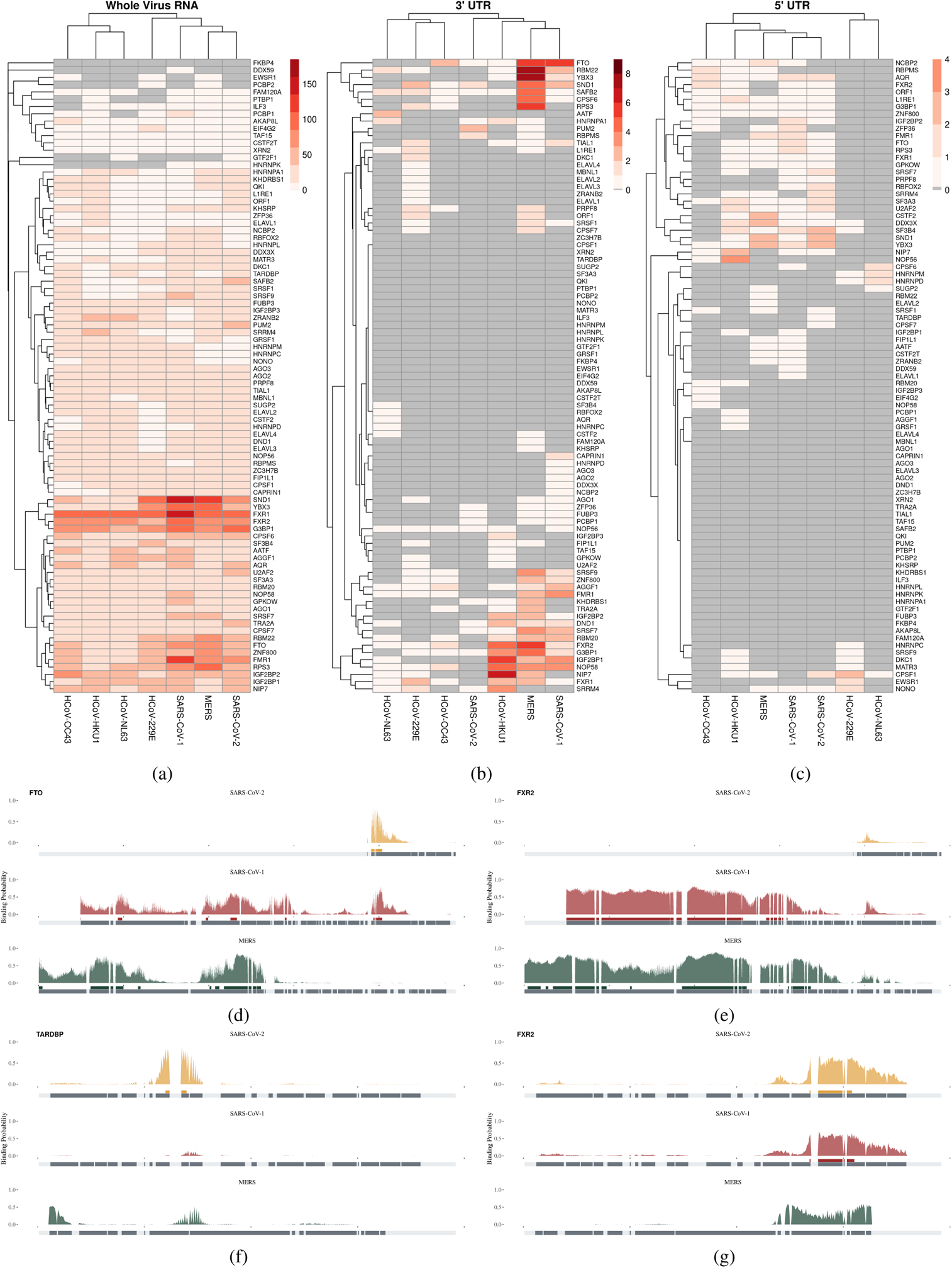
Comparison of SARS-CoV-2 and 6 other human coronaviruses. **a,b,c**. Binding sites were predicted over the seven human coronaviruses, and their number counted over the entire genome (**a**) or over the 3’ (**b**) and 5’ (**c**) UTRs. Hierarchical clustering was applied to evaluate the proximity between viruses in terms of binding sites composition. **d,e,f,g**. Examples of evolutionary conserved, gained, and lost binding sites between the three high-severity viruses MERS, SARS-CoV-1, and SARS-CoV2. Panel **d** shows an example for FTO binding sites found only in SARS-Cov-2 and SARS-CoV-1 in their 3’ UTRs. Panel **e** shows a binding site for FXR2 only shared between MERS and SARS-CoV-1 in their 3’ UTR. Panel **f** shows a binding site for TARDBP exclusive to SARS-CoV-2 in the 5’ UTR. Panel **g** shows a binding site for FXR2 only shared between SARS-CoV-2 and SARS-CoV-1 in the 5’ UTR.

#### Loss of FXR2-binding in SARS-CoV-2 3’ UTR

Figure 5e shows 3’ UTR binding of FXR2, a paralog of FMRP (fragile X mental retardation protein). Our model predicted extensive binding of FXR2 along the 3’ UTR of SARS-CoV-1 and MERS, while SARS-CoV-2 showed a complete lack of predicted FXR2 binding sites, owing to its significantly shorter 3’ UTR. On the other hand, Figure 5g shows that FXR2 binding is conserved in the 5’ UTR of SARS-CoV-1 and SARS-CoV-2. FMRP was previously shown to broadly bind along the entirety the 3’ UTR of the Zika virus (ZIKV) (74). However, while FMRP was suggested to act as a ZIKV restriction factor by blocking viral RNA translation, a significantly reduced ZIKV infection was observed upon knockdown of FXR2 (74).

#### Conserved FTO binding site in the 3’ UTR of SARS-CoV-1 and SARS-CoV-2

Altered expression levels of methyltransferase-like 3 (METTL3) and fat mass and obesity-associated protein (FTO) have been recently linked to viral replication (99). FTO is a demethylase (eraser) enzyme with enriched binding in the 3’ UTR of mRNAs in mammals (58). FTO has previously been suggested as a potential drug target against COVID-19 (97), as targeted knockdown has been shown to significantly decrease SARS-CoV-2 infection (99; 97; 6). Therefore, we investigated predicted binding of FTO to the 3’ UTR of SARS-CoV-2 and related viruses. Indeed, we observed that SARS-CoV-1, SARS-CoV-2 and MERS, as well as the less pathogenic viruses HCoV-HKU1 and HCoV-OC43 harbor at least one FTO binding site in their 3’ UTR (Figure 5b). Further, Figure 5d shows that while SARS-CoV-1 and MERS harbor multiple shared FTO binding sites along their 5’ UTR, SARS-CoV-2 only harbors one FTO binding site at the 3’ end of its 5’ UTR which is exclusively shared with SARS-CoV-1.

#### Newly acquired TARDBP binding in the SARS-Cov-2 5’ UTR

We next focus on TARDBP (also known as TDP-43) (Figure 5f), which was predicted to bind the 5’ UTR of a SARS-CoV-2 mutant in a recent study (60). TARDBP, a host protein implicated in pre-mRNA alternative splicing, has been shown to play a role in viral replication and pathogenesis in the context of coxsackievirus B3 infection (42). In contrast to the findings of Mukherjee et al. (60), our model identified a TARDBP binding site at the genomic range of 89-98 in the wild-type reference of SARS-CoV-2. Interestingly, in addition to observing a lack of predicted binding signal of TARDBP on the 5’ UTR of SARS-CoV-1 and MERS, we found a complete lack of TARDBP binding to the 5’ UTR of any of the other investigated coronaviruses (Figure 5c). This suggests that 5’ UTR TARDBP binding potential is newly acquired in SARS-CoV-2 and may affect its virulence.

### 2.5 A functional catalog of human RBPs with predicted SARS-CoV-2 interaction

To understand the functional impact of RBPs on the SARS-CoV-2-mediated COVID-19 disease, we set out to interrogate the breadth of publicly available OMIC research, thereby gathering supportive evidences for our 87 RBPs models (Figure 6). To this end, we collected 97 data sets of experimental research results from 22 studies (Methods 3.15) covering experimentally determined and predicted viral RNA - host RBP interactions as well as multi-level (OMICS) data related to SARS-CoV-2 cell line infections, shedding light on viral entry, protein-protein interactions and host cell regulation. Studies which are closer to disease phenotypes, like CRISPR cell survival assays and COVID-19 patient data, were also included. In addition, we collected evidence of direct involvement of RBPs in SARS-CoV-2 infection, as reported in the SIGNOR database, a manually curated resource of pathways and genes involved in SARS-CoV-2 (49). All data sets were harmonized and integrated through the use of knowing01 (kno) software to annotate RBPs by automated mapping of gene, variant and protein identifiers, yielding reported evidence of binding or regulation for 85 out of 87 (97.7%) RBPs models.

**Figure 6:**
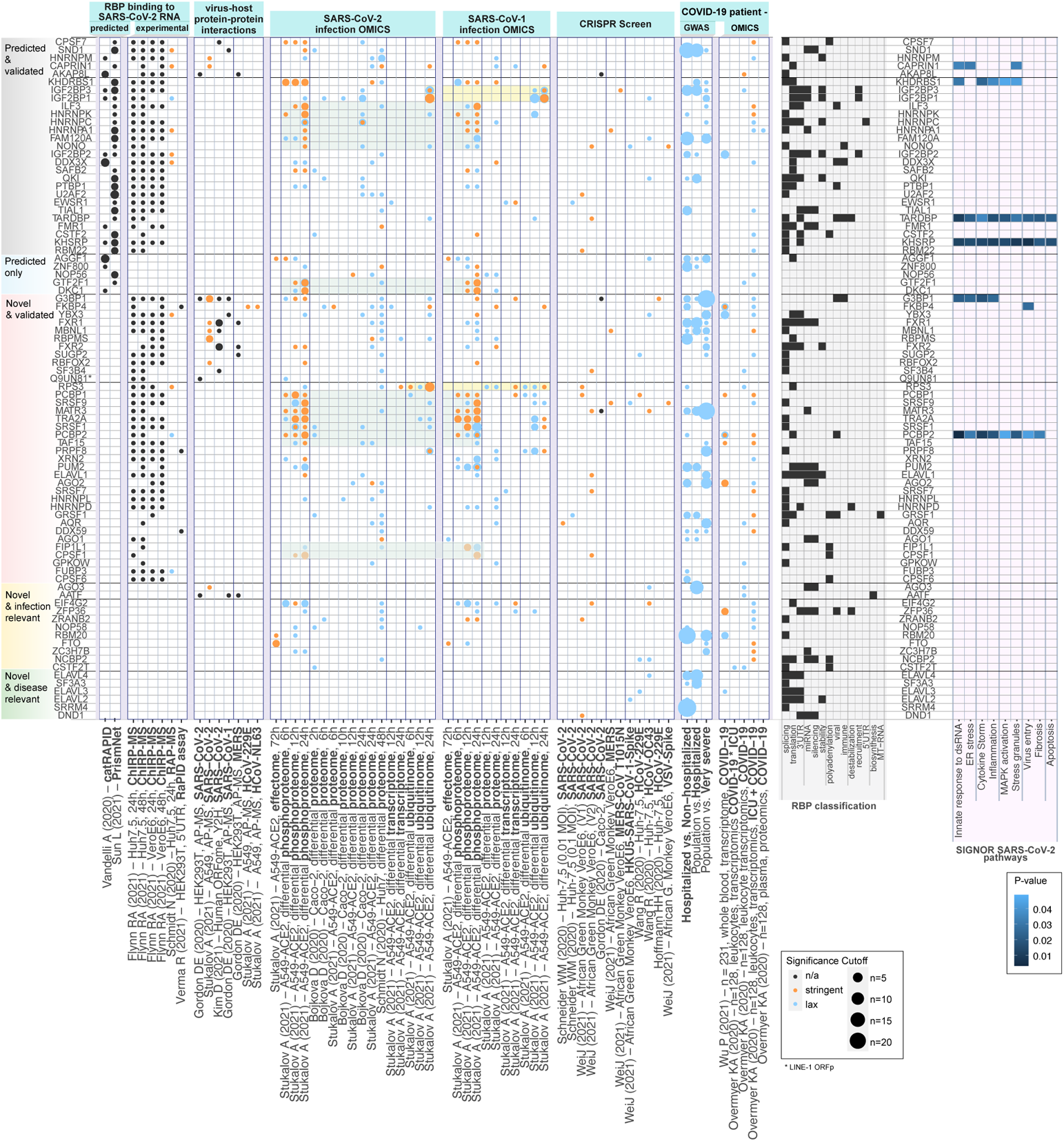
RBPs in context of public *in vitro* and patient OMICS data. RBP with model predictions (rows) annotated with experimental evidences found in 92 mulitOMIC publicly available research results (columns) followed by information from RBP classification and role in known SARS-CoV-2 pathways. From left to right: RBPs were manually assigned to five categories according to their annotation pattern. RBPs predicted to bind SARS-CoV-2 RNA by the other prediction methods catRAPID, PrismNET. RBPs binding to SARS-CoV-2 RNA determined experimentally by ChIRP-MS, RAP-MS and RaPID assay. Evidences of RBPs with stringent or lax significance cutoffs found in further 55 data sets across mulitple OMICS levels and experiment types were grouped by experimental context: Experimental viral-host protein interactions measured by AP-MS across various coronaviruses, SARS-CoV-2 and SARS-CoV-2 infection OMIC (timecourses), selected CRISPR studies, most recent GWAS data (release 6) by Host Genetics Initiative and blood-based patient OMICS data. Light green and yellow boxed highlight few patterns shared between SARS-CoV-2 and -1 infections. Classification of RBP according to their roles related to biological processes. Far right: Annotation of RBPs to pathways related to SARS-CoV-2 infections obtained from SIGNOR database.

We found that a large fraction (63 out of 87, 72.4%) of RBPs were identified to directly bind SARS-CoV-2 RNA using affinity-purification methods (69; 18) (Figure 6), validating the interaction of these RBPs with the viral RNA. Interestingly, only 32 out of 87 RBPs (36.8%) have previously had reported binding sites profiles over the SARS-CoV-2 genome by related methods catRAPID (87) or PRISMNet (80). We thus complement the knowledge on binding site locations over SARS-CoV-2 RNA with 55 RBPs uniquely explored by our framework, 36 of which are experimentally supported for viral RNA interactions (labeled as ‘NOVEL validated’, Figure 6). Our holistic comparison revealed that the majority of explored RBPs (75, 86.2%) were previously reported to be part of host-pathogen PPI networks and cellular pathways which are altered during infection by either SARS-CoV-2, SARS-CoV-1 or both (Figure 6). In addition, 34 out of the 87 (39.1%) were identified as essential genes in CRISPR knock-out screenings, highlighting the importance of RBPs in the infection process, immune response and viral replication, through direct interaction with the viral genome. Although no RBP co-localizes with loci associated to COVID-19 severe disease courses (GWAS) under genome-wide significance, we identified 44 (50.6%) RBPs with nominal significance. When considering the total of 2,730 coding genes co-localizing nominally associated loci, this represents a significant enrichment for RBPs (odds ratio of 7.8, Fisher test p-value <2.2e-16), suggesting their importance in patient’s course. Finally, a small set of our predicted-binding RBPs was shown to be supported only from CRISPR screens or found deregulated across COVID-19 patients, without evidence of viral RNA binding from previous studies, neither functional evidence in molecular networks altered by SARS-CoV-2 infection (labeled as ‘NOVEL & disease relevant’, Figure 6). Taken together, the large overlap between the RBPs we selected and the different resources considered confirms that hijacking host RBPs is crucial to the infection life cycle of the virus, through the direct binding of these RBPs to the viral genome only or in combination with host-pathogen protein-protein interactions.

## 3 Material and Methods

### 3.1 ENCODE data and preprocessing

Enhanced CLIP (eCLIP) datasets were obtained from the ENCODE project database, which comprises 223 eCLIP experiments of 150 RBPs across two cell lines, HepG2 and K562. For RBPs with experiments in both cell lines, we selected only data of eCLIP experiments from the HepG2 cell line for downstream analysis, as those were demonstrated to yield higher performing models (compared to K562) in previous studies (5). Narrow peaks of each eCLIP library were taken directly from ENCODE and preprocessing was performed as follows: for each of the two replicates of a given eCLIP experiment, peaks were first intersected with mRNA locations obtained from the GENCODE database (Release 35) and only overlapping peaks were retained. Next, the 5’-end of each peak was defined as the cross-linked site, as it usually corresponds to the highest accumulation of reverse transcription truncation events. A 400bp window was then centered around the cross-linked site for each peak, defining the input window of the downstream model. Input windows of both replicates were intersected reciprocally with a required overlap fraction of 0.75, ensuring that only those peaks which are present in both replicates are considered for downstream training set construction. Finally, the top most 50,000 windows with a read-start count FC of 2.0 above the control (SMInput) experiment were selected for each RBP.

### 3.2 Pysster training set construction

For each RBP, a classification dataset of bound (positive) and unbound (negative) RNA sequences was constructed. Positive samples were obtained by taking corresponding 400nt peak-region windows from the previous step (3.1), while two distinct sets of negative samples were generated. First, 400nt long regions which did not overlap with binding sites of the given RBP were sampled from transcripts harboring at least one binding site. This constraint ensures that the transcript is expressed in the experimental cell type and would not be observed as RBP-binding in other cell types. The second set of negative samples was generated by randomly sampling binding sites of other RBPs. This ensures that any CLIP-seq biases (such as U-bias during UV-C cross-linking (79), (93)) are present in both positive and negative samples and prevents the model from performing a biases-based sample discrimination during the training. Together, this yields a three-class training set, where class 1 corresponds to positive samples and class 2 and 3 correspond to negative samples. Samples of class 2 and 3 were sampled at a 3:1 ratio with respect to class 1. Finally, generated samples were randomly split into train, validation and test sets at a ratio of 70:15:15, respectively.

### 3.3 Pysster model

The *pysster* Python library (5) was used for implementation of the model which consists of three subsequent one-dimensional convolutional layers, each with 150 filters of size 18, followed by a single fully connected layer with 100 units. The ReLU activation function is applied to each intermediate layer output and a maximum pooling layer is added after every convolutional layer. Finally, a fully connected layer with 3 units, one for each of the three output classes, is added. Dropout (75) with a rate of 0.25 was applied to each layer, except for input and output layers. The model was trained with the Adam optimizer (41) using a batch size of 512 and a learning rate of 0.001. For each RBP, we trained for at most 500 epochs and stopped training in case the validation loss did not improve within the last 10 epochs.

### 3.4 Pysster binary classification threshold

As pysster models are trained as a 3-class classification problem with class imbalance, we re-calibrate each model for the binary classification task by introducing a binary decision threshold *t_m_* on the predicted positive-class probability scores. For each model *m*, *t_m_* is defined as the threshold which maximizes the F1 performance (Section 3.7.1) of the model with respect to bound vs. unbound binary classification obtained by pooling class 2 and 3 samples into a common ‘unbound’ class. This threshold is used to identify bound regions in the viral sequence (Section 3.9).

### 3.5 DeepRiPe model

We obtained pre-trained DeepRiPe models from Ghanbari et al. (23) and retained models for 33 out of the 59 RBPs, filtering out models where no informative sequence motif could be learned by the model. The PAR-CLIP-based models used in this study are modified versions of the DeepRiPe neural network, where only the sequence module to extract features from the RNA sequence is used. Briefly, the model consists of two convolutional layers, one fully connected layer and one output layer that contains *k* sigmoid neurons to predict the probability of binding, one for each RBP. Each convolutional layer has a rectified linear unit (ReLU) activation, followed by a max pool layer and a dropout layer with probability of 0.25. 90 filters with length 7 and 100 filters of length 5 for the first and second convolution layers, respectively. The fully connected layer has 250 hidden units and a ReLU activation. Details in data preparation and model training are outlined in Ghanbari et al. (23).

### 3.6 Single-nucleotide predictions

The pysster and DeepRiPe positive-class prediction score corresponds to the probability that input RNA sequence is bound by the RBP of interest. By design, this score is assigned to the entire input sequence, although RBP binding sites are much more local, usually spanning only a few nucleotides (14). To obtain single-nucleotide binding site probabilities from both pysster and DeepRiPe models along an RNA sequence, we employ a one-step sliding-window approach to scan over a given RNA sequence, where the predicted positive-class probability score is assigned to the center nucleotide of the input window. In order to obtain predictions over the entire RNA sequence, the 5’ and 3’ sequence ends are 0-padded.

### 3.7 Pysster performance evaluation and model selection

#### 3.7.1 Precision-recall and F1 performance

As the validation loss was monitored for the purpose of early-stopping, the precision-recall (PR) and F1-score performance of the pysster models was evaluated on the test set. Models with an area under the PR curve (auPRC) of less than or equal to 0.6 were deemed poor quality and thus excluded from the downstream analysis.

#### 3.7.2 Performance in practice

Training datasets are sampled at a fixed positive-negative ratio which hardly reflects the ratio of bound and unbound sites of RNA transcripts found *in vivo*. In practice we expect that for some transcripts regions, binding sites of a particular RBP are not observed over several kilo-bases, while other regions, such as 5’ and 3’ untranslated regions (UTRs), might harbor a dense clustering of binding sites. To measure the ability of pysster models to accurately predict *de novo* RBP binding-sites along whole-length RNA transcripts, we introduce the concept of Performance-In-Practice (PIP), which measures how well the single-nucleotide prediction score of the model correlates with binding sites identified by eCLIP. For a given RNA sequence, the PIP of a model is defined as the Spearman correlation coefficient (SCC) between the truncated prediction scores 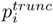 and a binary vector obtained by labeling all positions that fall within eCLIP binding sites with 1 and 0 otherwise. Here, 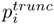 refers to a modified version of the prediction score *p_i_* defined as

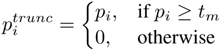

where *t_m_* is a threshold obtained for each model as outlined in Section 3.4. For each model, we perform extensive PIP analysis on the human transcriptome as follows. First, we select the set of transcripts which contain at least one binding site for it. From this set, we uniformly draw 100 transcripts without replacement as hold-out transcripts. Subsequently, we intersect positive and negative training samples with the hold-out transcripts and discard all samples that overlap with any of the hold-out transcripts before retraining pysster on the remaining training samples. We use the resulting models to predict along the hold-out transcripts as described in Section 3.6 and compute the PIP score for each hold-out transcript. Finally, models with a median PIP score of less than or equal to 0.1 were excluded from downstream analysis.

### 3.8 Estimating significance of prediction scores

To directly control the false positive rate of binding site prediction from both pysster and DeepRiPe models on the viral genome, we estimate prediction score significance via an RNA sequence permutation test. In order to obtain a null-distribution of predictions (positive-class) scores, we first compute the di-nucleotide frequencies on the viral RNA. Next, we perform frequency-weighted sampling of di-nucleotides to construct a set of *N* = 10, 000 null-distributed inputs. Null-distributed prediction scores for each model are then obtained by predicting on those sequences. A p-value is assigned to each observed prediction score *p_i_* in the viral sequence by computing the fraction of scores from the null distribution 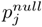 for which 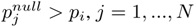.

### 3.9 Identifying RBP binding sites

We identify RBP binding sites on the viral RNA sequence using predicted single-nucleotide binding probabilities (Section 3.6) together with estimated p-values (Section 3.8). For each pysster model, we classify nucleotides in the viral RNA as “bound” if the predicted probability score is equal or greater than the estimated binary threshold *t_m_* (Section 3.4) and the score is found to be significant (*p <* 0.01). Regions with a consecutive stretch of bound nucleotides of at least length 2 are then defined as a RBP binding site. Neighboring binding sites that are spaced by less than 10 nucleotides are merged to a single binding site. Note that for DeepRiPe models, nucleotides in the viral RNA are considered “bound” if the probability score is found to be significant (*p <* 0.01) and no score threshold is applied.

### 3.10 Base-wise feature attribution via Integrated Gradients

To gain insight into which RNA sub-sequences are driving factors for RBP binding, we compute sequence importance scores using Integrated Gradients (IGs) (81; 23). Starting from an input baseline, IG performs a step-wise linear path interpolation between the baseline and the actual input sequence and computes the gradients of the interpolated inputs with respect to an output neuron. That is, we obtain a vector of importance scores over the input sequence which indicate which nucleotides of the input contributed most toward the prediction. Here, we choose the 0-vector (i.e. the one-hot encoding of all nucleotides is set to 0) as the baseline and perform 50 baseline-input interpolation steps. To obtain sequence importance scores for a given binding site, we compute IGs with respect to an input window centered around the binding site. For sequence-motif construction, the heights of nucleotides in the input sequence is given by the feature attribution weights.

### 3.11 Analyzing mutations in variants of concern

Variant information of 11 SARS-CoV-2 viral variants (alpha, beta, delta, epsilon, eta, gamma, iota, kappa, lambda, mu, omicron) was obtained from the UCSC genome-browser for the SARS-CoV-2 virus (17), and converted into VCF format. For each strain, we first created a ‘mutated’ strain-specific genome, using the viral reference sequence and the set of strain-defining variants. We then center a window at the reference position of each genomic variant and extract the mutated sequence for subsequent prediction via each model. We note that for cases were genomic variants are in close proximity with each other, extracted windows might contain multiple mutations. This is crucial, as only the combination of multiple variants might lead to gain or loss of RBP binding. The resulting prediction score on each alternative allele (ALT) is then compared with the prediction score of the same window on the reference sequence (REF). To quantify the impact of each mutation, we compute a *delta* score between the prediction score of ALT and REF sequence:

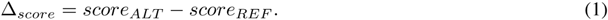

Mutations with a positive delta score sign represent ‘gain-of-binding’ (GOB) events, while mutations with negative sign represent ‘loss-of-binding’ (LOB) events. To further narrow down the set of mutations, we compile a subset of mutation that lead to a gain-or loss-of-binding (GOB and LOB), defined as instances where (in case of LOB) the REF score is passing the binding site score threshold and p-value (Sections 3.4 and 3.8) while the ALT does not, or vice versa (in case of gain of binding). As for binding site calling (Section 3.9), we use a significance level of 0.01 as p-value threshold for both pysster and DeepRiPe models.

### 3.12 *In silico* mutagenesis

We perform *in silico* probing of the effects of all possible point-mutations on RBP binding across the SARS-CoV-2 genome. At each viral genome position, the reference base was mutated to each of the three alternative bases. Subsequently, prediction was performed on the input windows derived from each ALT allele using all high-quality pysster models. Finally, as described in Section 3.11, an impact score is computed and a set of change-of-binding mutations is compiled.

### 3.13 Comparative analysis of human coronaviruses

Besides SARS-CoV-2, we obtained reference sequences for 6 other human coronaviruses, including SARS-CoV-1, MERS, HCoV-229E, HCoV-HKU1, HCoV-NL63 and HCoV-OC43 from NCBI (68). Using high-quality models from both pysster and DeepRiPe (Section 3.7), we perform single-nucleotide binding prediction along each viral RNAs (Section 3.6). Next we compute prediction empirical p-values for each viral sequence (Section 3.8) by generating a dedicated null distribution of scores for each virus and RBP. RBP binding sites across viruses were then identified as described in Section 3.9. We evaluate genomic-element preference across a subset of shared viral genomic locations (ORF1ab, E, N, M, S, 5’ UTR, 3’ UTR) for each RBP and virus by intersecting the predicted set of binding sites of each virus with its RefSeq annotations. To compute multiple sequence alignments (MSA) between genomic elements of betacoronaviruses, we use the ClustalO (72) algorithm with default parameters.

### 3.14 Functional annotation of RBPs

To assess the potential role of RBPs with predicted binding on viral RNA sequences, we manually curated all RNA-related functions of the 87 RBPs with good predictive models using the GeneCards, Uniprot and RBP2GO databases (77).

### 3.15 Public COVID-19/coronaviruses OMICS data

To assess regulatory information of RBPs across available coronavirus/COVID-19 multiOMICS data, we downloaded evidence from 22 studies. We imported study-relevant supplementary tables via knowing01, which harmonizes data tables and links results to molecular information, like human gene symbols, UniProt identifier, variant positions as available in the proprietary CellMap unified data model (Version 2022/03). A list of 87 RBPs with good model performance were loaded as list of Gene Symbols. To ensure that all RBP human gene symbols are identically named in African Green Monkey OMICS data, we used VeroE6 cells linked to human symbols.

A total of 97 research results were grouped into the following study types:

- extended interactomes from experimental determined of host RBP-SARS-CoV-2 interactions using affinity purification and mass spectrometry (18; 69; 89)
- computational predictions of host RBPs-SARS-CoV-2 interactions in the 5’ UTR, 3’ UTR and Spike S genomic region of the viral RNA with either catRAPIDomics (87) or the PRISMNet tool (80)
- viral-host protein-protein interactions (PPIs) measured by affinity-purification followed by mass spectrometry (24; 78)] and yeast two hybrid screenings (40)
- multiOMICS data, including the regulation of the host proteomics, phosphoproteomics, ubiquitinomics and transcriptomics up to 24 hours after coronavirus infection (4; 78), as well as the effectome, which includes deregulated host proteins 72 hours after SARS-CoV-2 induced expression of each of the viral proteins (78)
- CRISPR phenotype screens probing cell survival few days after viral infection with single genes knockouts in human (24; 30; 70; 91) or African green monkey [(92)] cell lines
- genome-wide association studies (GWAS) linking human genetic variation to COVID-19 disease severity (64)
- patient OMICS data, including proteomics and transcriptomics regulation of whole blood, serum or plasma of mostly inpatients (10; 12; 13; 22; 57; 63; 71; 95)

To filter for significant regulation in each data set, we applied significance cutoffs per study result. We chose to select two different significance levels to get lists of regulation with a stringent (adjusted p-value < 0.01) and a lax (adjusted p-value < 0.1) cutoff threshold, whenever available. Few data sets only provided raw p-values for which we used with lower cutoffs. Patient transcriptomics data were used with much lower cutoffs, due to the inflation of regulated genes on typical cutoffs. For GWAS data we employed a genome-wide (p-value < 5e-08) and nominal (p-value < 0.01) significance cutoff, for stringent and lax cutoffs, respectively. Finally we annotated all 87 RPBs with regulated molecules via the knowing01 Annotate feature and visualized the number of evidences of RBPs in each data set in a count matrix.

## 4 Discussion

Strong evidence suggests that human RBPs are critical host factors for viral infection by SARS-CoV-2, yet there is no feasible experimental approach to map exact binding sites of RBPs across the SARS-CoV-2 genome systematically. To combat this knowledge gap, we constructed the first *in silico* human-virus RBP-RNA interaction map for SARS-CoV-2 using predictions from pysster (5) and DeepRiPe (23) models trained on a large cohort of eCLIP and PAR-CLIP datasets, respectively. The use of high-capacity CNN classifiers represents a significant improvement over previous computational studies performing motif scanning over the SARS-CoV-2 genome (75; 3), as it enables the learning of more complex binding syntax and thus the detection of binding sites for RBPs with no cleanly defined sequence motif. This is evident by the fact that we observed high performance for RBPs without annotations of binding motifs in literature. On the other hand, we demonstrated that deep learning methods are by no means black boxes, as we recovered known binding motifs for several RBPs (including QKI, RBFOX2 and TARDBP) using gradient-based attribution methods. Together with stringent performance evaluation and conservative selection of high-quality models, these results suggest that our predictions represent bona fide binding sites. In a recent study, the PRISMNet deep learing model was used to infer binding of 42 host RBPs to the SARS-CoV-2 genome (80). However, predicted binding sites by PRISMNet are restricted to the 5’ and 3’ viral UTR regions are rather large, with some spanning over hundreds of nucleotides, while RBP binding usually only occurs across short stretches of RNA *in vivo*. In contrast, our approach generated single-nucleotide binding probabilities across the entire viral genome and may therefore yield a more complete picture of the binding landscape of human RBPs to SARS-CoV-2.

Our study identified known, as well as novel human RBPs to interact with SARS-CoV-2 (Figure 6). Further, the generated binding map provides a rich resource for future functional studies, in particular for investigating the role of the SARS-CoV-2 protein-RNA interactome in context of the viral life cycle. For instance, binding site predictions may be used to accelerate the discovery of host RBPs that engage in both pro-and anti-viral functions by directly interacting with the viral RNA. Further, predictions may aid in the identification of functional sites on the viral RNA that can be therapeutically targeted by RNA drugs, such as anti-sense oligonucleotides, to interfere with host RBP binding. In addition to constructing a RBP binding map on the SARS-CoV-2 reference sequence, we quantified the impact of sequence variant from 11 SARS-CoV-2 strains, including the alpha, delta and omicron viral strains.

Additionally, we applied a systematic *in silico* mutagenesis of all positions in the SARS-CoV-2 genome, pinpointing mutations associated with particularly high impact, which could represent potential high-risk variants to monitor in the future. Our analyses confirmed that our models can effectively be used to identify mutations with high-impact potential using the prediction scores, either for mutations observed in viral variants of concern (Figure 4a) or from *in silico* mutagenesis (Supplementary Figure 3). Such mutations can be evaluated further through the computation of attribution maps, highlighting important nucleotide in a given window of interest, and how their importance is impacted by the mutation. In previous studies variants of concerns have been prioritized through their potential impact on the sequence of viral proteins, in particular the Spike protein. Our results complement these findings, and enable to better understand the efficiency of specific lineages of SARS-CoV-2 in the context of RBP-viral RNA interactions, providing with a map of mutations of high potential for hijacking important host RBPs, or on the contrary evade binding of anti-viral RBPs. With our comparative analysis of RBP-RNA interactions across seven coronaviruses we contribute to the identification of genomic features and factors which confer unique characteristics to SARS-CoV-2 transmission and pathogenicity, compared to SARS-CoV-1, MERS, and less pathogenic coronaviruses. Both variants of concern and comparative analysis highlight gain-or loss-of-binding affecting host RBP-viral interactions and therefore pinpoint RBPs which can be prioritized for further screening.

We integrated knowledge of our predicted RBPs across other pathogens, host-viral protein-protein interactions, numerous studies collecting functional and phenotypic data, such as GWAS and CRISPR screens, as well as multi-omics COVID-19 patient data, in order to pinpoint RBPs with clinical significance. By this analysis, we mainly identify five sets of RBPs predicted to interact with the SARS-CoV-2 genome. The first set comprises core RBP predictions with numerous independent evidences in the scientific literature of their involvement in regulation of viral infection, included SARS-CoV-2. Proteins in this core set are confirmed by additional *in silico* methods, as well as experimental assays to bind SARS-CoV-2, and identified as deregulated or affected in multi-omics studies and/or CRISPR, GWAS and patient data of SARS-CoV-2 infection. Among them, we find several known regulators of viral processes, such as the hnRNPR viral restriction factors (65), the IGF2BP1-3 RBPs, which are mainly ubiquitinated during SARS-CoV-2 infection (78) and linked, through GWAS, to poor disease outcome (34), as well as key regulators of SARS-CoV-2 infections such as the stress granules-associates RBPs CAPRIN1 and KHDRBS1 (37), associated to pathways such as ER stress, Inflammation, cytokine storm and others (Supplementary Table 3), the pro-viral DDX3X factor (9) and the host factor NONO (65), previously shown to promote innate immune activation in HIV infection (44). Important regulators of mRNA splicing (QKI, PTBP1 and U2AF2), and other processes (TARDBP, TIAL1) are also part of this group of RBPs. Notably, many of the RBPs we highlighted throughout our binding site analysis on the SARS-Cov-2 genome, impacts from mutations in viral variants, or comparative genomic changes of binding sites fall into this group. For instance, TARDBP and QKI are two RBPs that are well supported, in particular through experimental identification of their binding to the viral RNA, in addition to OMICs support and CRISPR (TARDBP) or GWAS (QKI). We also identify TARDBP as a particularly important RBP in the context of SARS-CoV-2 infection due to the prediction of a unique binding site in the virus 5’ UTR, when compared to SARS-CoV-1, MERS and other coronaviruses. A second set of RBPs comprises 36 proteins uniquely predicted by our framework as binders of SARS-CoV-2, which harbour experimental extensive support.

An example of RBP of interest in this group is the Serine/arginine-rich splicing factor 7 (SRSF7). Previous studies have shown that SRSF7 interacts with coronavirus RNA (76). It has also been suggested that this spliceosome protein could be sequestrated by the viral genome, the later thus acting as a sponge through these putative binding sites, to alter host splicing processes. Among the high-impact mutations in the SRSF7 gene position 23,604 (S protein gene) is found mutated across multiple strains, with different alternative nucleotides: a C>A transversion is found in alpha and mu variants, while a C>G transversion is found in delta and kappa variants. Both mutations are associated to a positive delta score, therefore a gain of binding. This position has been suggested by previous studies to be a major driver of the increased infection efficiency of these viral variant, as a modifier of the S protein sequence (P680R) (52), although additional studies indicate that other mutations may be required for an actual effect ((53; 98)). The gain of binding we identify here could also suggest that the translation of the S gene into the protein is improved through the recognition of the newly created binding site by SRSF7.

Besides SRSF7, the large number of binding sites for splicing factors at the 5’ UTR of the SARS-CoV-2 (cluster 6, Figure 3c) and the pervasive binding of several host and viral restriction factors (cluster 4, Figure 3c) suggests that these RBPs are likely to get sponged on the viral genome and by that modulate post-transcriptional regulatory networks in the host cell.

One other interesting RBP in this group is represented by FXR2, paralog of FXR1 and FMR1 which are identified as direct binders of SARS-CoV-2 (Figure 6). Recent evidence suggests that FXR2 selectively interact with MERS viral proteins but not with viral proteins from SARS-CoV-1 and SARS-CoV-2 [(24)]. While we find evidence of FXR2 binding along the SARS-CoV-2 genome, this is in agreement with the results of our comparative analysis with other human coronaviruses, where we observe extensive binding of FXR2 along the 3’ UTR of SARS-CoV-1 and MERS, but depletion of FXR2 bindidng in the SARS-CoV-2 3’ UTR. Together with the evidence of genetic association of FXR2 to COVID-19 disease severity (35) our findings suggest a fine-tuning role of FXR2 in regulating the severity of the infection.

From these two sets, we can also highlight many RBPs with functions related to endoplasmic reticulum processes. SARS-Cov-2 utilizes the endoplasmic reticulum (ER)-derived double membrane vesicles (DMVs) as replication centers. RNA viruses, included SARS-CoV-2, contains several instances of an RNA regulatory motif, called SECReTE motif (27) which facilitates localization to the ER and increases viral protein translation, as well as viral replication. Such motif is also found in some human mRNAs encoding for proteins involved in innate immunity and associated with epithelial layers targeted by SARS-CoV-2. This suggests that host and pathogen might compete for ER-associated RBPs and this might make the host more vulnerable to the infection. Among our validated RBPs in set 1 and 2 (Figure 6) we identified several SECReTE-associated RBPs, defined as those proteins where more than one fourth of their predicted binding sites overlapped instances of the SECReTE motif on the SARS-CoV-2 genome. These include FUBP3, KHSRP and MATR3, already identified previously as important host or restriction factors for other RNA virus infections (65). Interestingly, we linked MATR3 to several CRISPR studies showing that this factor is essential for SASR-CoV-2 replication, as well as to many nominal variants in all GWAS data (Figure 6). MATR3 physically interacts with G3BP1, another predicted RBP in this set which been found to interact specifically with SARS-CoV-2 nucleocapsid (N) protein, control viral replication and localize (together with MATR3) at stress granules where G3BP3 is taken away from its typical interactions partners (62). Our and previous data (Figure 3) suggest that direct binding of G3BP3 and MATR3 to the SARS-CoV-2 RNA could constitute an additional mechanism used by the virus to interfere with the G3BP3-MATR3 PPI network and impair stress granule formation. The fact that G3BP3 binding is enriched in correspondence of the gene encoding for protein N (Figure 3c) might also suggest a direct regulation of this transcript by this RBP in a sort of feedback loop manner.

The other three sets of RBPs predicted to bind SARS-CoV-2 correspond to 1) proteins with *in silico* support from other predictive tools, but no experimental validation of direct binding to SARS-CoV-2 (named ‘Predicted only’); 2) novel candidate SARS-CoV-2 binders, uniquely predicted by our method, no experimental validation but large functional support from host-pathogen PPI, CRISPR and patient omics data (named ‘Novel infection relevant’), and 3) putative novel regulators that lack so far functional evidence across studies but were nonetheless found to be deregulated in COVID-19 patients (named ‘Novel disease relevant’). The fat mass and obesity-associated protein (FTO) is an example of a newly identified regulatory RBP for SARS-CoV-2. FTO is a demethylase (19), and while it has been suggested that the virus could hijack the host epigenome [(2)], a recent study showed that the viral genome itself was methylated (51), with a negative effect on viral replication efficiency. Besides the predicted binding pattern, FTO also presented numerous important gain-or loss-of-binding across many viral strains. Although there was no clear trend towards systematic loss of binding of FTO across the viral variants, we were able to point out multiple close-by mutations in the alpha variant that were associated to a significant loss, around the position 28,280 (Figure 4b). Finally, the FTO protein was identified as key risk factor for obesity, which is also a known risk for COVID-19 severity. FTO coding region harbored also nominal genetic associations to COVID-19 severity (variant lowest p-value 0.0053). Interestingly, FTO was additionally found to be significantly regulated on gene level in blood serum of patients admitted to ICU care (adj. p-value 7.72E-06) (63). A small set of novel predicted RBPs, with little to no experimental evidence across multiple functional studies, includes the ELAVL2-4 factors, the DND1 RBP and the splicing factors SRRM4 and SF3A3 (Figure 6). Interestingly, ELAVL2-4 RBPs, found in our analysis to be SECReTE motif-associated RBPs, and SRRM4 RBP are neuron-specific proteins and were found, through our integrative analysis, to be deregulated in COVID-19 patients. This points to novel promising candidates whose molecular mechanisms can be further investigated experimentally.

## 5 Conclusion

Viruses depend on essential host factors at all stages of their infection cycle. One family of host factors, RNA-binding proteins (RBPs), are involved in multiple aspects of post-transcriptional regulation and are characterized by their ability to bind to short RNA motifs. While several RBPs have been associated with SARS-CoV-2, some of which may represent drug-able targets for anti-viral therapy, cost and time constraints render a comprehensive experimental profiling of human RBPs to the SARS-CoV-2 RNA infeasible. To fill this knowledge gap, we instead identified binding of human RBPs to the SARS-CoV-2 genome computationally. Here, we used the pysster and DeepRiPe frameworks together with data from over 200 eCLIP and PAR-CLIP experiments to train RBP binding site predictors on the basis of convolutional neural networks (CNN). By applying stringent performance filters, we obtained a set of high-quality prediction models for 88 RBPs and created an *in silico* binding map of human RBPs along the SARS-CoV-2 genome at single-nucleotide resolution. Predicted binding profiles of RBPs suggested that groups of RBPs exhibit similar binding patterns on the viral genome and that RBPs within these group may be functionally related, for example, by being associated to the SECReTE motif important for efficient viral replication. We identify RBPs with clinical relevance, by analyzing our data in the context of functional and clinical studies, including genetic screens and COVID-19 patient data. We further utilized trained models to score the impact of strain-defining sequence variants across 11 SARS-CoV-2 strains. Several variants that result gain or loss of RBP-binding were identified, some of which simultaneously impact the binding of multiple RBPs or which are conserved in multiple viral strain. In addition to the analysis of observed variants, we quantified the impact of hypothetical variants by performing extensive *in silico* mutagenesis, generating all possible point mutations across the SARS-CoV-2 genome. We believe that this resource will greatly aid researchers in assessing the impact of newly identified viral variants. Finally, we predicted RBP-binding across 6 other human coronaviruses (including SARS-CoV-1 and MERS) and identified several conserved binding sites as well newly acquired binding sites in SARS-CoV-2.

All generated results, including fully trained models, predicted binding sites across SARS-CoV-2 and other coronaviruses, variant impact scores across 11 viral strains and impact scores of hypothetical variants are publicly available at https://sc2rbpmap.helmholtz-muenchen.de/. We believe that our results give new insight into the role of RNA-binding proteins in context of SARS-CoV-2 infection and represents a rich resource for further research on how SARS-CoV-2 hijacks the host cell’s RNA regulatory machinery for viral replication and evasion of immune response.

## Code and Data Availability

Training data and pre-trained models, together with scripts for training and prediction are available at https://github.com/mhorlacher/sc2rbpmap. RBP binding sites on the SARS-CoV-2 genome and variant impact scores for 11 viral strains are available at https://sc2rbpmap.helmholtz-munich.de.

## Acknowledgments

This work was supported by the Helmholtz Association under the joint research school “Munich School for Data Science - MUDS to M.H., S.O., G.C., P.S. and A.M., the Deutsche Forschungsgemeinschaft (SFB/TR501 84 TP C01) to A.M. and L.M., the Helmholtz Association AeroHEALTH grant to A.M. and Y.H., additionally supported by the Joachim Herz Foundation for Y.H. MG and UO are supported by the Berlin Center of Machine Learning (BZML) funded by the German Ministry for Education and Research (BMBF).

## Author Contributions

**Marc Horlacher:** Conceptualization; Data pre-processing and curation; Machine learning model training and prediction; comparative genomics analysis; viral strains analysis; Interpretation of results; Visualisation; Methodology; Implementation of the Dashboard; Writing – original draft; Writing – review & editing. **Svitlana Oleshko:** Conceptualization; Data curation; RBP map clustering and visualisation; SECReTE motif and viral strains analysis; Interpretation of results; Writing – review & editing. **Yue Hu:** Conceptualization; Data curation; model predictions; downstream statistical analysis; viral strains analysis; Interpretation of results; Methodology; Writing – review & editing. **Mahsa Ghanbari:** Analysis of PARCLIP data and machine learning model training; Writing – review & editing. **Giulia Cantini:** Implementation of the Dashboard. **Patrick Schinke:** Implementation of the Dashboard. **Ernesto Elorduy Vergara:** Conceptualization; Methodology. **Florian Bittner:** Software engineering for public data integration and analysis. **Nikola S. Mueller:** Conceptualization; Supervision; Public data curation and analysis; Visualisation; Writing – original draft; Writing – review & editing. **Uwe Ohler:** Conceptualization; Supervision; Funding acquisition and resources. **Lamber Moyon:** Conceptualization; Supervision; Viral strain analysis; Interpretation of the results; Visualisation; Methodology; Writing – original draft; Writing – review & editing. **Annalisa Marsico:** Conceptualization; Supervision; Methodology; Visualisation; Interpretation of the results; Funding acquisition and resources; Writing – original draft; Writing – review & editing.

## 8 Supplementary Tables

**Supplementary Table 1:** Comparison of high quality pysster and DeepRiPe models

| RBP | cor of<br>pred score | cor of<br>p-value | # common<br>binding sites |
| --- | --- | --- | --- |
| TARDBP | 0.640832 | 0.40309 | 6 |
| CSTF2 | 0.459011 | 0.21621 | 6 |
| IGF2BP1 | 0.387823 | 0.39983 | 9 |
| PUM2 | 0.383309 | 0.35263 | 10 |
| CSTF2T | 0.331395 | 0.22239 | 3 |
| QKI | 0.279760 | 0.14371 | 5 |
| IGF2BP2 | 0.171838 | 0.21092 | 7 |
| IGF2BP3 | 0.073798 | 0.05951 | 5 |
| CPSF6 | 0.153344 | 0.26078 | 2 |
| FXR1 | 0.012354 | 0.14136 | 8 |
| FXR2 | 0.080191 | 0.19433 | 5 |
| EWSR1 | 0.009787 | 0.06610 | 0 |

**Supplementary Table 2:**
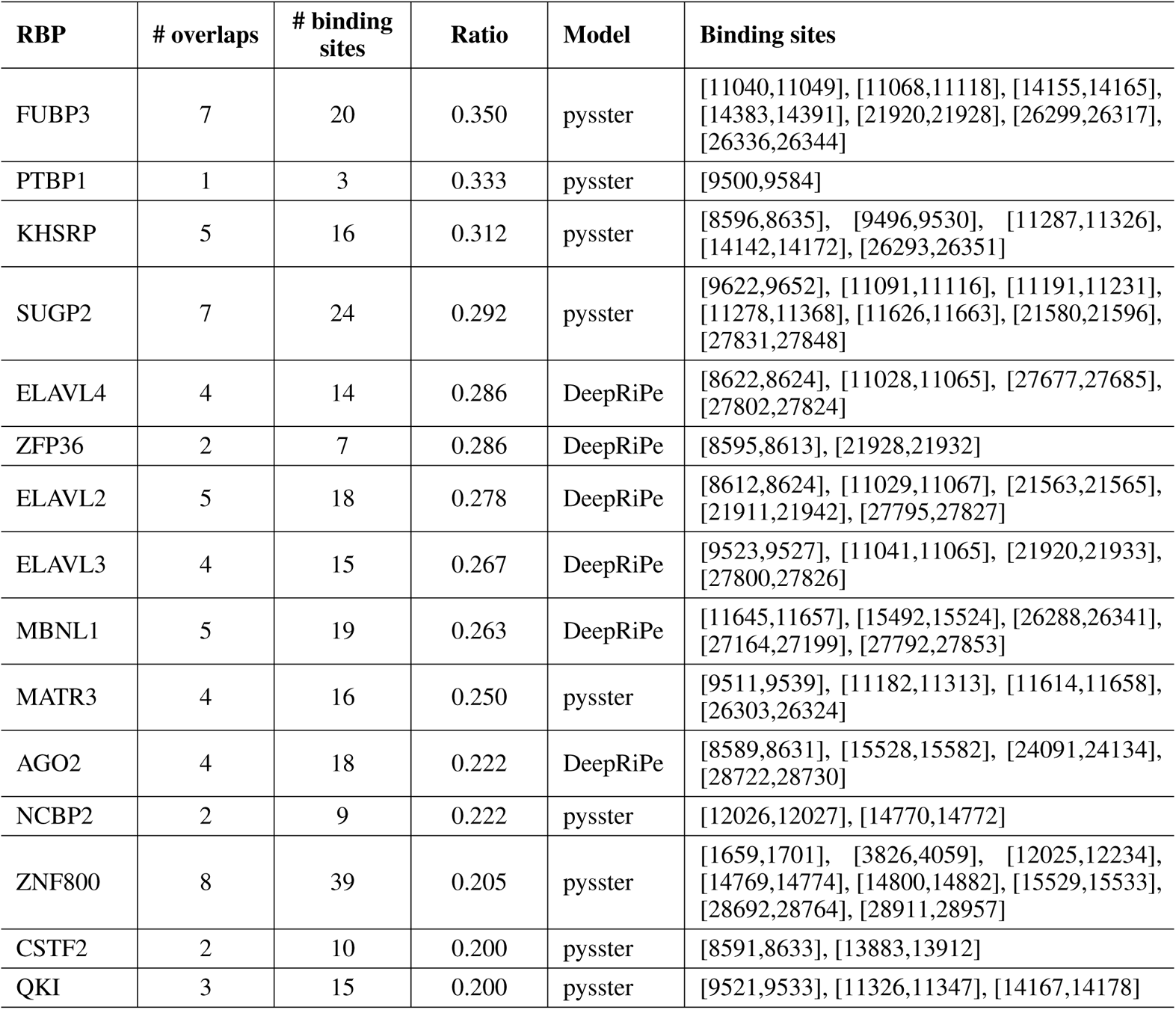
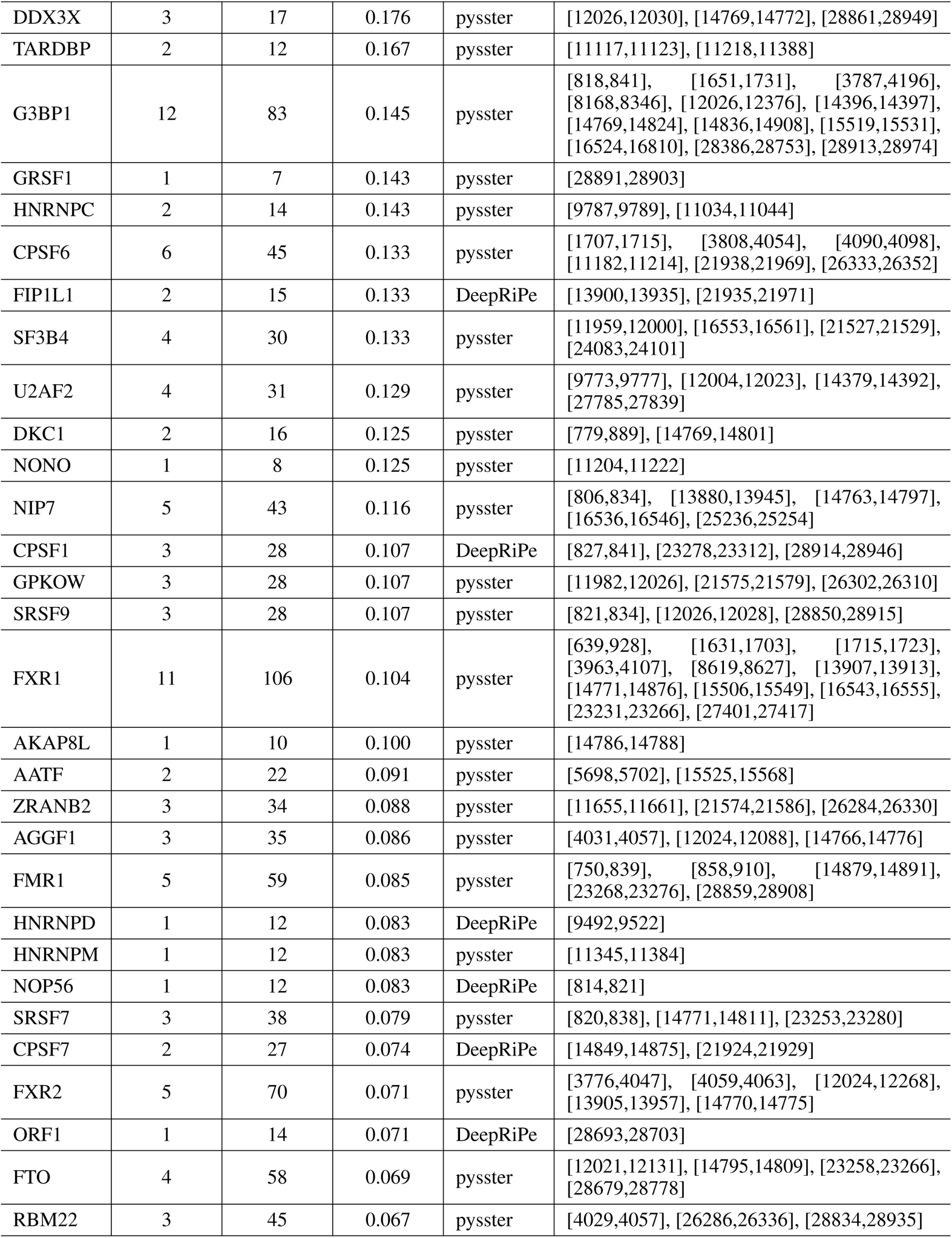
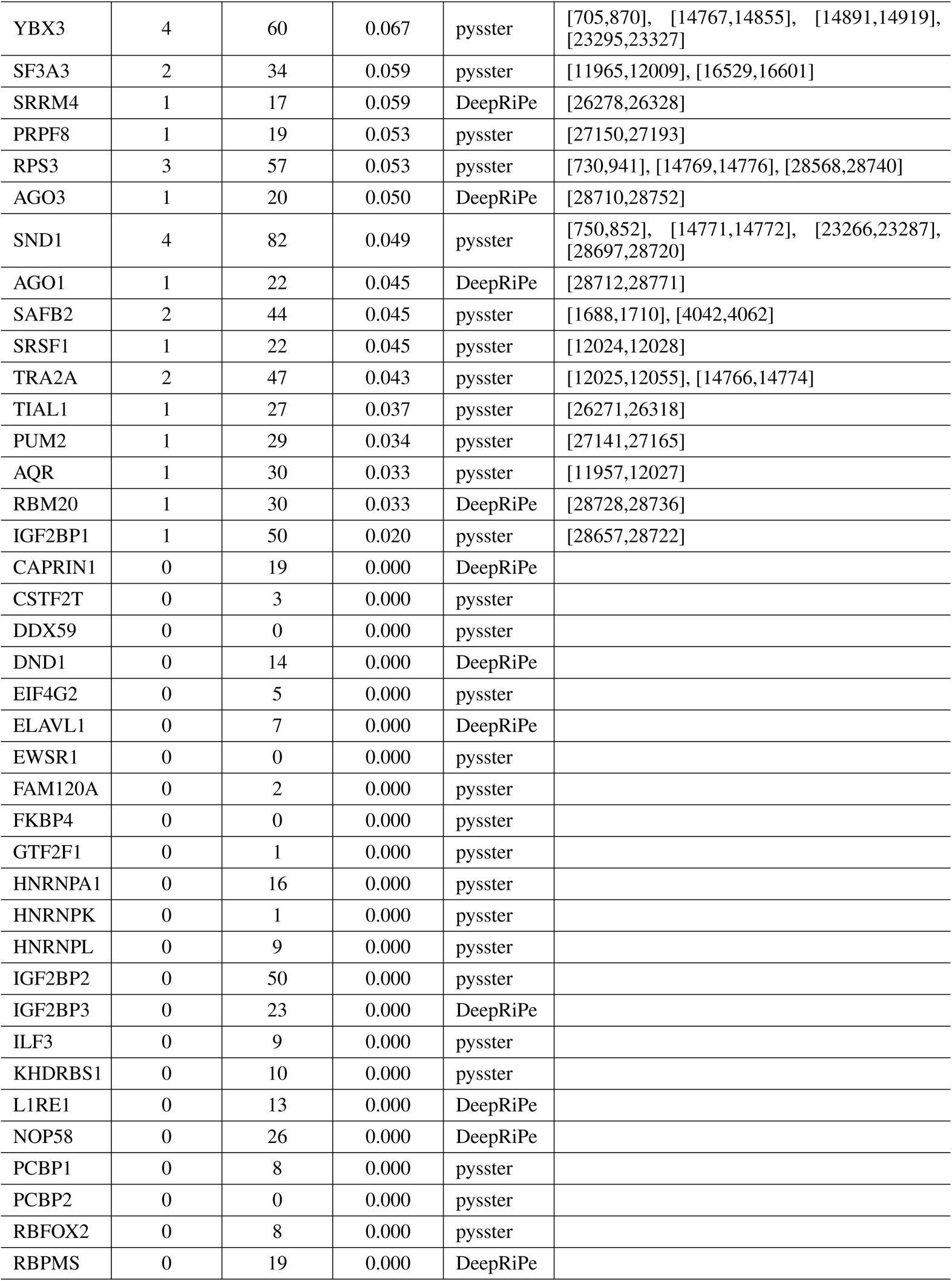
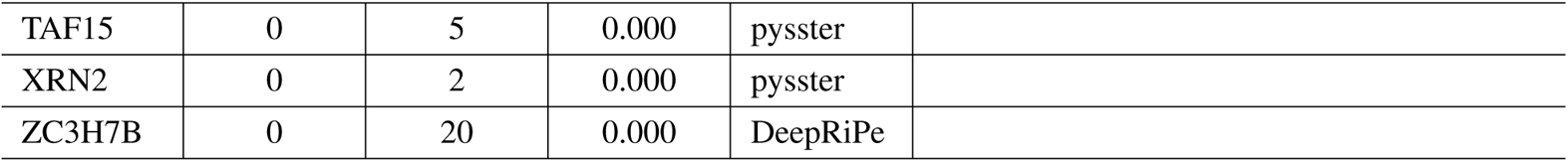
Overlap of pysster and DeepRiPe binding sites with SECReTE motif

**Supplementary Table 3:**
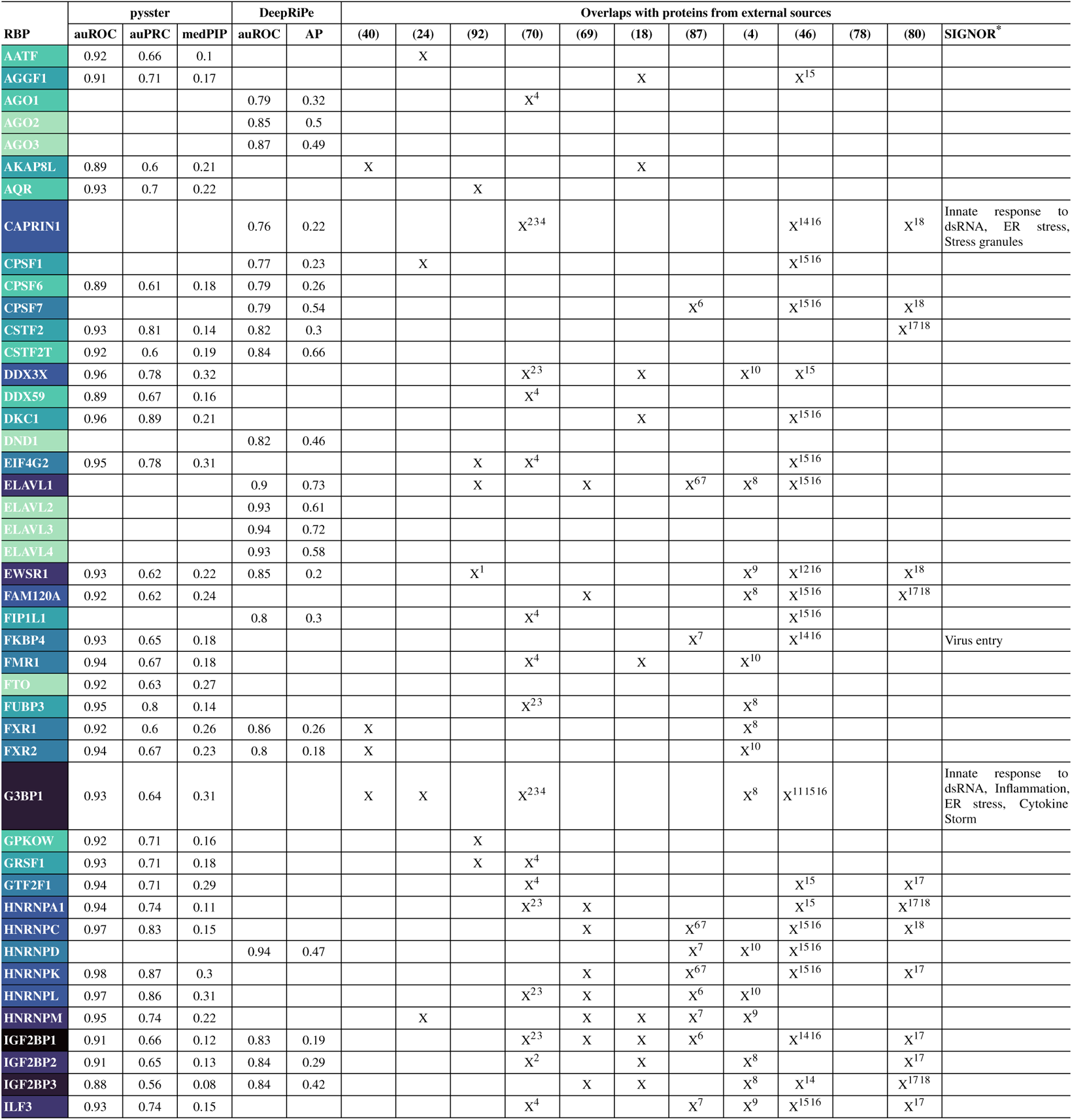
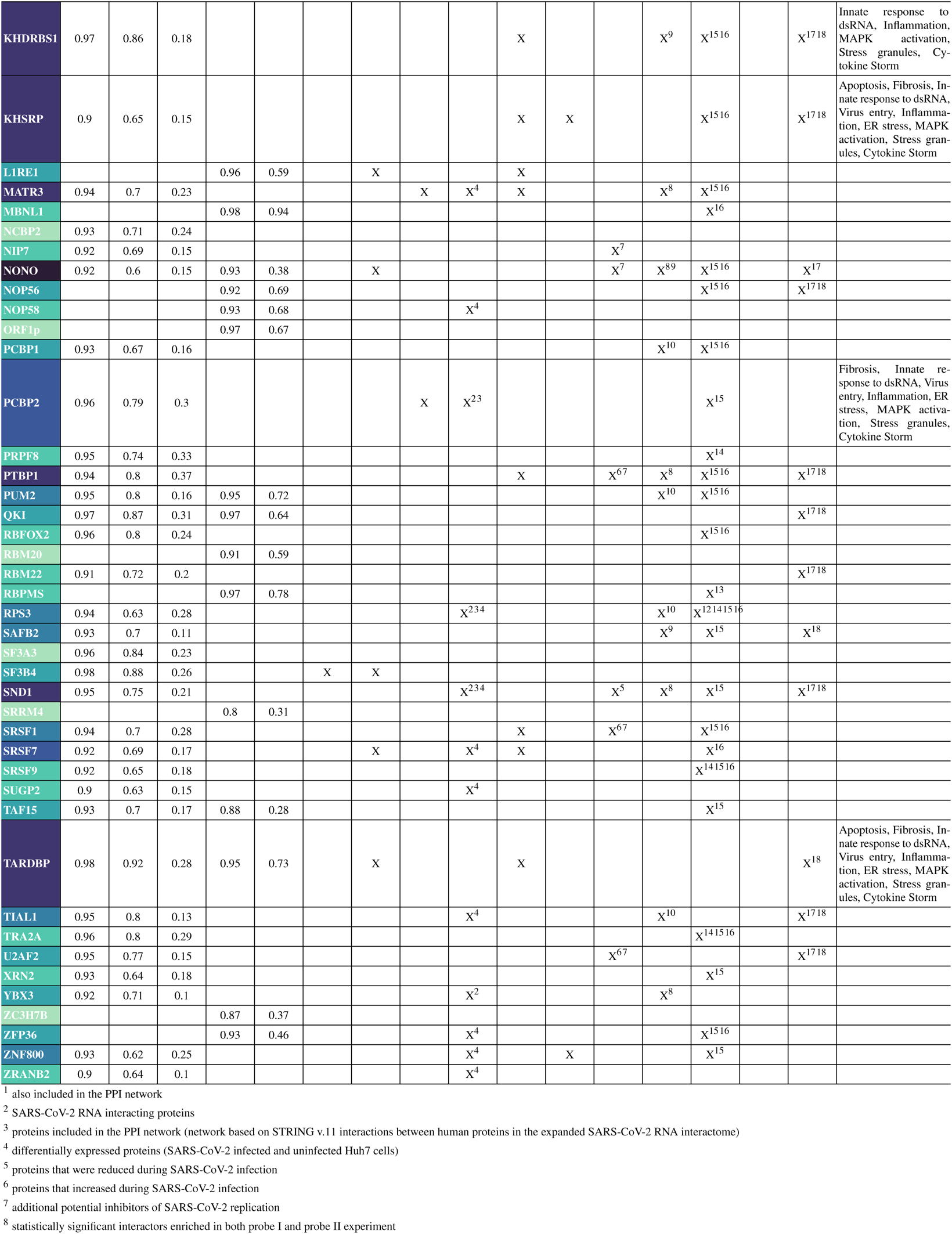
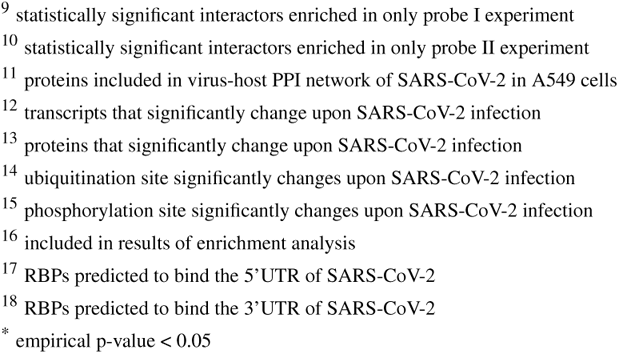
Overlap of pysster and DeepRiPe models with proteins from external sources

## 9 Supplementary Figures

**Supplementary Figure 1:**
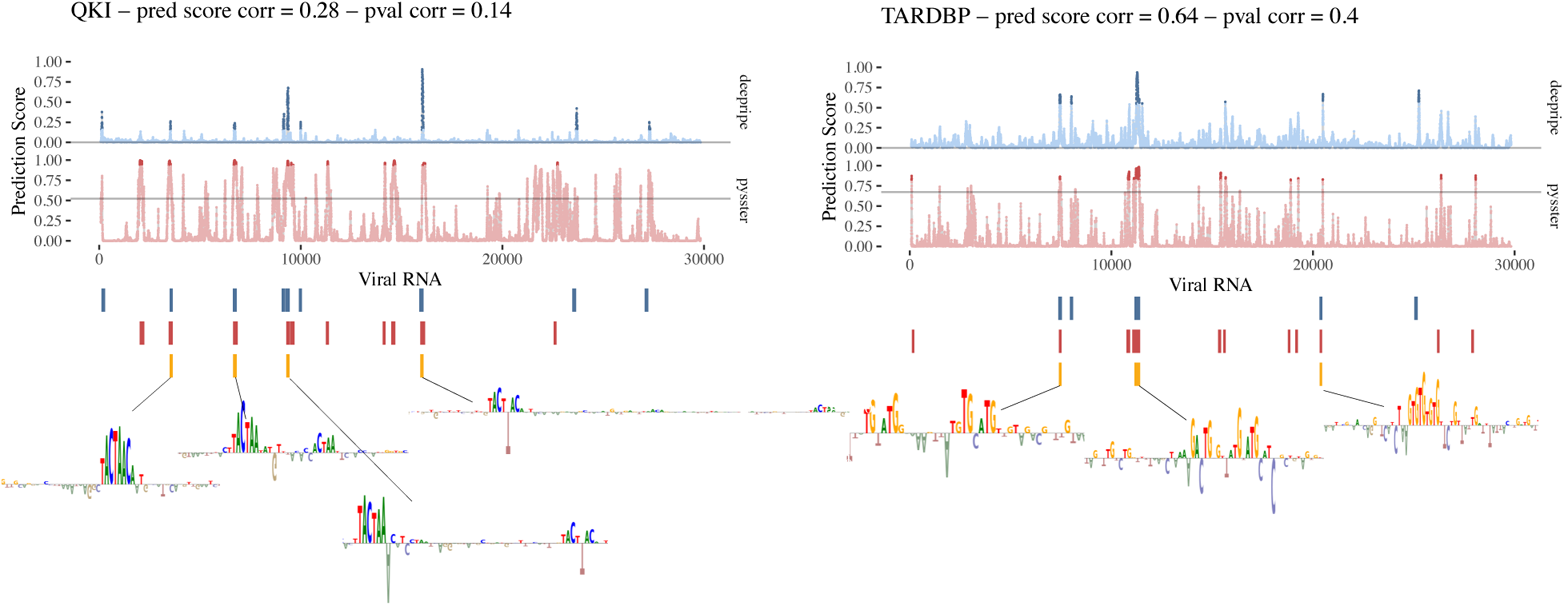
RBP binding pattern on the SARS-CoV-2 genome between the two methods, pysster and DeepRiPe. Comparison of single-nucleotide probability scores of binding for two RBPs, QKI (left panel) and TARDBP (right panel). Significant binding sites, commonly predicted by both methods are shown underneath the probability plots together with their corresponding learnt motifs from the attribution maps. Prediction score correlation and p-value correlation given in the header.

**Supplementary Figure 2:**
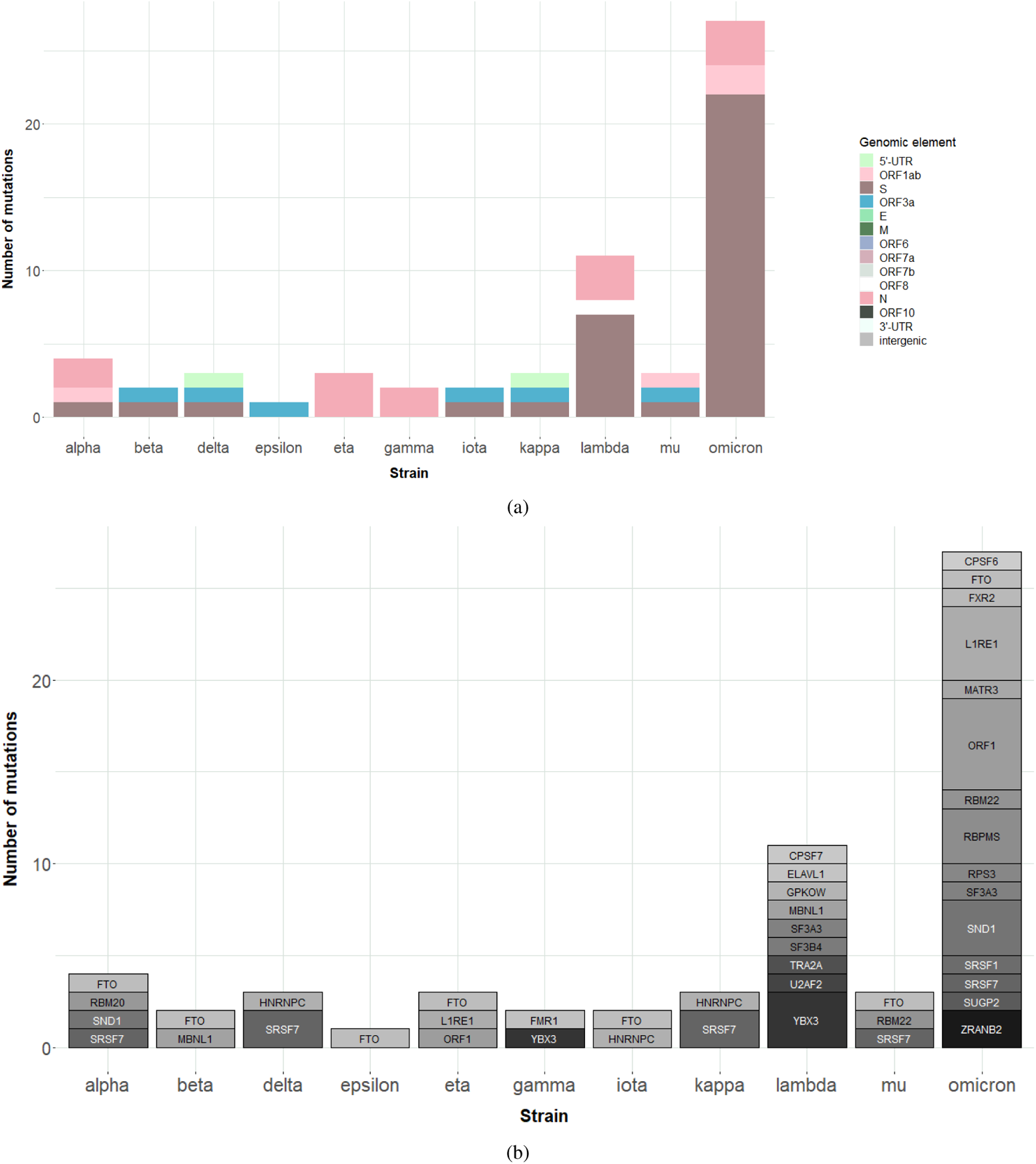
Impact of variants of concern on predicted binding sites. **a**. Accumulation of high-impact variants of concern in viral components for each lineage. The subset of high-impact variants here corresponds to the one represented in Figure 4a, i.e. the top 20% of binding-impacting variants. **b**. Accumulation of impacted RBP sites for each lineage. The same subset as in (a) was used here.

**Supplementary Figure 3:**
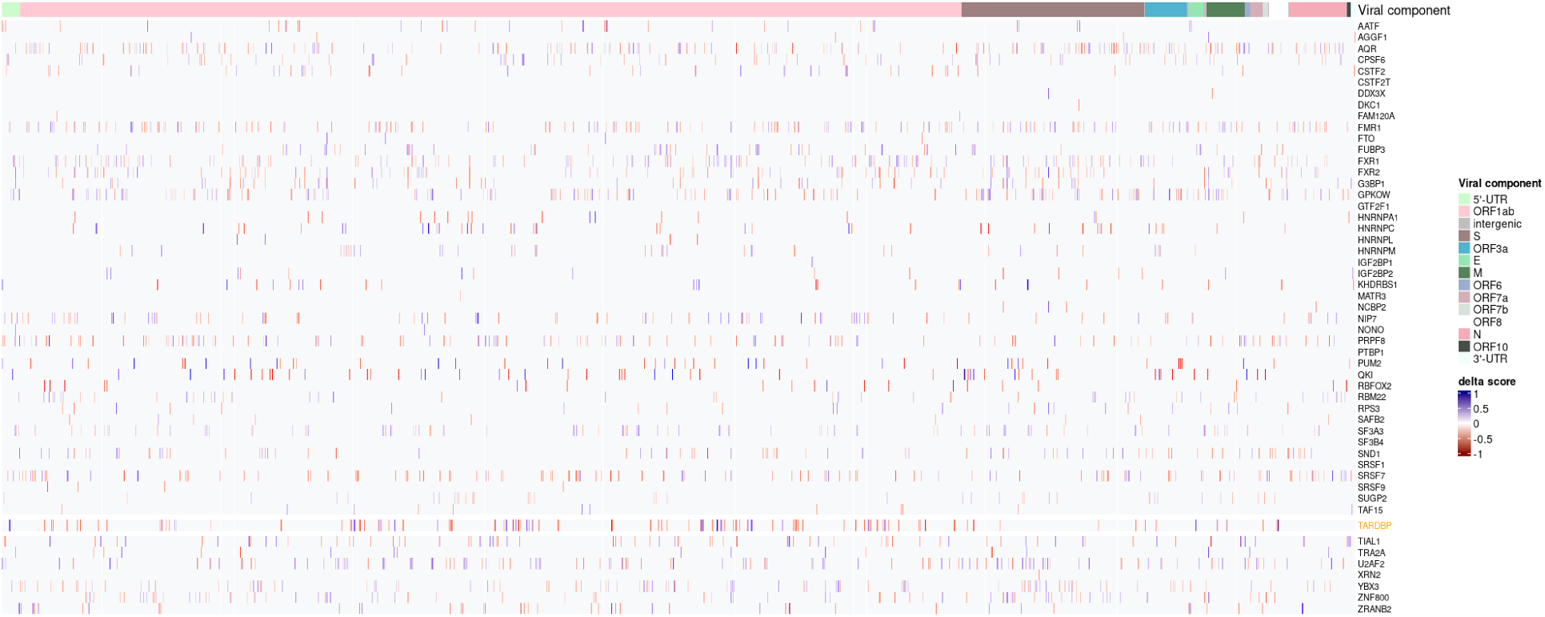
*In silico* perturbation analysis of SARS-CoV-2. Nucleotides across the viral genome were perturbed towards the three alternative bases and the alternative base with resulting the highest delta score considered for downstream analysis. Here, we show the delta score heatmap across positions with at least one gain- or loss-of-binding event across all RBPs.

**Supplementary Figure 4:**
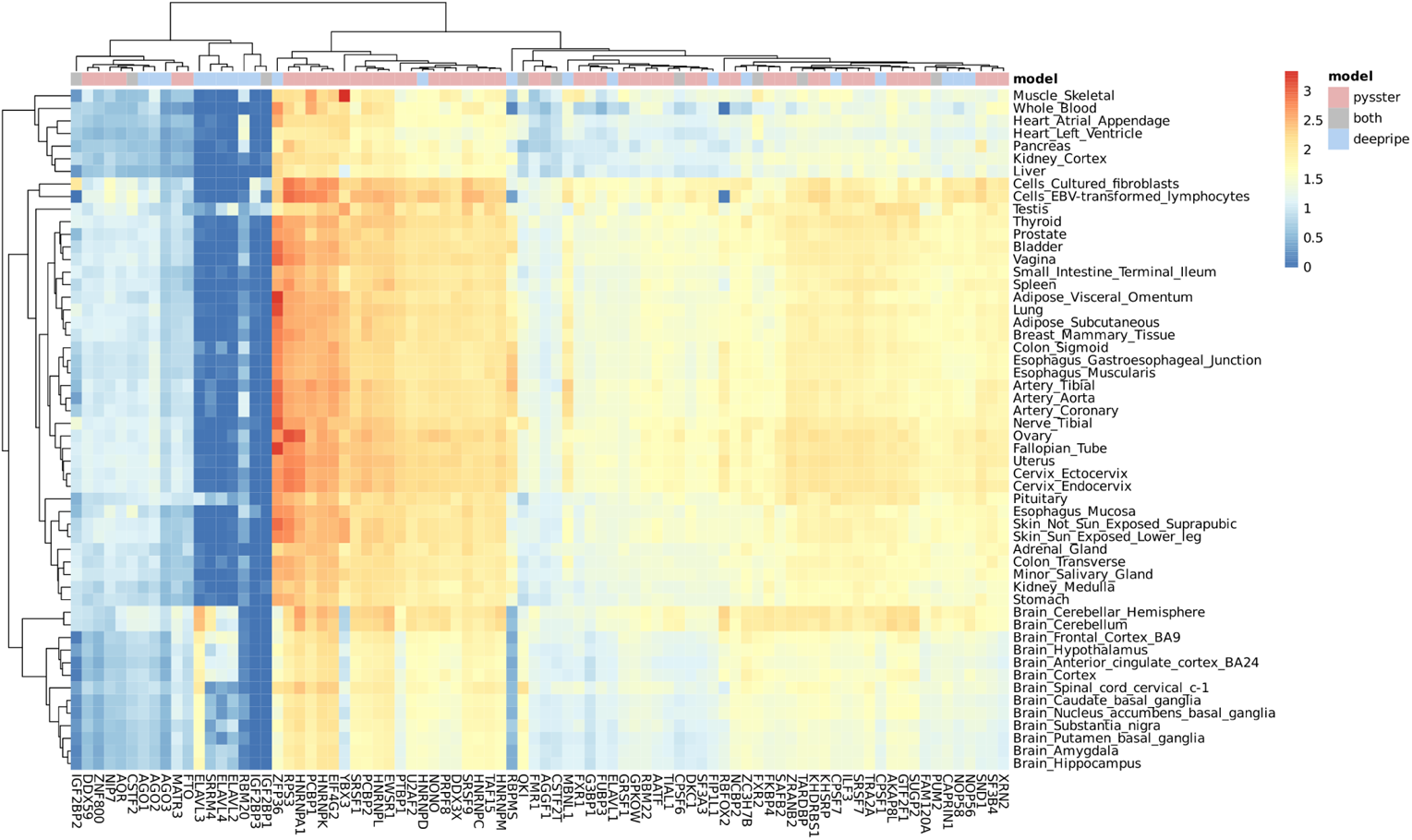
Expression of RBPs in tissues across the body: Median expression values in log10 transcript per million (TPM) of RBPs across 54 sub-tissue types from the Genotype-Tissue Expression (GTEx) project (7). RBPs from different methods color coded above the heatmap: pysster-exclusive in red, DeepRiPe-exclusive in blue, and shared between models in grey.

